# Generating human parvalbumin interneurons through 3D glia reprogramming

**DOI:** 10.1101/2024.09.06.609855

**Authors:** Christina-Anastasia Stamouli, Alexander Degener, Efrain Cepeda-Prado, Andreas Bruzelius, Emil Andersson, Jessica Giacomoni, Aurélie Delphine Vorgeat, Srisaiyini Kidnapillai, Oxana Klementieva, Malin Parmar, Victor Olariu, Daniella Rylander Ottosson

## Abstract

Parvalbumin (PV) interneurons are crucial for synaptic plasticity, and their damage or loss is linked to various neurological disorders. Yet, generating these cells of human source *in vitro* is challenging, limiting advancements in cell repair and disease modelling. We introduce a novel approach to derive human PV neurons through direct reprogramming of glial precursor cells (GPCs). Using ectopic expression of GABAergic neuronal genes, GPCs efficiently convert into GABAergic interneurons in 3D culture environment within weeks and achieve functional neuronal maturity. Single-nuclei RNA sequencing identified a distinct PV neuronal cluster with high maturity and characteristics of PV chandelier subclass that are equivalent to bona fide human interneurons. Trajectory analysis revealed a distinct glia-to-PV interneuron conversion pathway, involving several new transitory genes, with potential for functional importance for PV derivation. Our data introduces a new strategy for generating human PV interneurons, promising significant implications for future generation of patient-specific PV neurons both *in vitro* and *in vivo*.

**Highlights:**

- A novel approach to derive human PV interneurons by direct glia reprogramming.
- First comprehensive transcriptomic profiling of induced human PV interneurons.
- Induced PV interneurons are of chandelier subtype with transcriptional similarity to *bona fide* interneurons.
- Successful glia-to-PV interneuron conversion passes through a specific reprogramming trajectory and involves key genes with functional potential.

## Introduction

In the cerebral cortex, excitatory principal glutamatergic neurons and inhibitory GABAergic interneurons are the two main neuronal types responsible for excitation and inhibition^1^. Among these, the subtype of parvalbumin (PV)-expressing interneurons have unique morphological and functional properties that makes it crucial in neural circuits and memory processing^2^. Recent evidence shows that PV interneurons are often lost or dysfunctional in several neurological and neuropsychiatric diseases and states, such as epilepsy and schizophrenia^3–7^. This loss or dysfunction is not just a symptom but a critical step in the underlying neuronal network dysfunction and cognitive decline, underscoring the need for improved strategies for generating and replacing PV interneurons. Experimental studies indicate that cell restoration of GABAergic interneurons via cell transplantation could counteract disease pathology^5,7–9^. However, generating subtype-specific PV interneurons *in vitro* from stem cell or fetal sources has proven to be difficult^9–11^. This is possibly due to the limited time for neuronal maturation in 2D cultures, the unmet need for complex culture systems for synaptic integration and the high energy demands of PV interneurons.

Direct neuronal reprogramming offers a promising alternative approach to stem cell differentiation strategies for cell replacement therapies. Here, non-neuronal cells can be converted into neurons using specific combinations of key transcription factors, miRNAs, and/or cocktails of chemical compounds^12^. Importantly, this approach bypasses the pluripotency step, making it feasible for in-brain repair^13^. Using direct reprogramming, we and others have shown that mouse resident glial cells can convert into subtype-specific neurons^14–20^, including PV interneurons, in the living mouse brain^21–23^. Glial progenitor cells (GPCs) are particularly suitable for *in vivo* reprogramming due to their proliferative nature in adulthood and widespread distribution throughout the brain parenchyma^24–26^. However, transferring this technique to the human system has been challenging due to the late embryonic development of human GPCs^27^.

Herein, we utilized a stem cell differentiation protocol to derive human GPCs (hGPCs) for direct neuronal reprogramming^28,29^. The hGPCs were transduced with a combination of GABAergic fate determinants^30,31^ and reprogrammed in a recently established 3D setting that promotes long-term cell viability and synaptic connectivity^32^. In 21 days, the hGPCs rapidly reduced their glia protein and gene expression and simultaneously induced the expression of neuronal genes. Single-cell transcriptomic analysis revealed characteristic pallial interneuron genes and a distinct PV-enriched cluster in only 13 days resembling *bona fide* PV interneurons. Importantly, lineage trajectory analysis of the reprogramming process showed a specific PV reprogramming pathway involving several novel dynamic genes. *In vitro* manipulation of these can potentially hold functional importance for successful PV derivation both from stem cells and through direct neuronal reprogramming.

In conclusion, this study presents a novel approach for rapidly deriving human PV interneurons via direct reprogramming, with potential applications for future *in vivo* cell replacement therapies and patient-derived *in vitro* models.

## Materials and methods

### hESC-derived GPCs production

hGPCs were obtained from hESCs (RC17 Roslin Cells, cat.no. hPSCreg RCe021-A, p26–30). hESCs were cultured on tissue plates coated with LN521 (0.5 μg/cm^2^, Biolamina, Sundbyberg, Sweden) and kept in IPS-Brew XF medium (StemMACS, Miltenyi, Bergisch Gladbach, Germany). Weekly passages were performed using EDTA (0.5 mM, Gibco, Thermo Fisher, Waltham, MA, USA). hESCs were differentiated into hGPCs according to established protocols ^28^.

### Lentiviral production

Previously described vectors^30^ were used for the direct conversion of hGPCs. Briefly, we used a mix of constitutively expressed and doxycycline-dependent transcription factors. The constitutively regulated transcription factors were open reading frames (ORFs) of *Ascl1*, *DLX5* and *LHX6*. The doxycycline-regulated transcription factors were ORFs of *Sox2* and *FOXG1*. All the transcription factors were in third-generation lentiviral vectors (LVs). hGPCs were always co-transduced with LV containing the FUW-M2rtTA plasmid (#20342, Addgene, Watertown, MA, USA) for the expression of doxycycline-regulated transcription factors. The lentiviral vector used to overexpress *RORA*, (pLV [Exp]-EGFP-TRE>hRORA with vector ID VB230913-1017xen) was constructed and packaged by VectorBuilder. LVs were produced as previously described^33^ and titrated with quantitative PCR analysis^34^. The range of the titers was 2.5×10^8^ to 3.42×10^9^.

### Conversion of hGPCs and generation of converted spheroids

After 2-3 weeks of thawing, hGPCs were detached with a cell scraper and dissociated into single cells with Accutase (StemPro, Thermo Fisher; Waltham, MA, USA). 100.000 single hGPCs were mixed with the lentiviral reprogramming cocktail [multiplicity of infection (MOI) of 1–2 per vector] in 60 μl of media and seeded in 96-well U-bottom plates (#CLS7007, Corning, NY, USA) for self-aggregation. After 24 hours, 60 μl of fresh medium containing doxycycline (5 μg/ml, Duchefa, Haarlem, Netherlands) were exchanged to induce tet-regulated transgene expression. Three days after transduction, media was replaced by neural differentiation (ND) medium, which consisted of NDiff227 (Takara-Clontech, Gothenburg, Sweden),with doxycycline (5 μg/ml for 3D and 2 μg/ml for 2D), small molecules (CHIR99021[2 mM, Axon, Groningen, Netherlands]; SB-431542[10 mM, Axon, Groningen, Netherlands]; noggin[0.5 mg/ml, R&D Systems R&D Systems, Minneapolis, MN, USA]; LDN-193189[0.5 mM, Axon, Groningen, Netherlands]; VPA[1 mM, Merck Millipore, Burlington, MA, USA]) and growth factors (LM-22A4[2 mM, R&D Systems, Minneapolis, MN, USA]; GDNF,[2 ng/ml[R&D Systems, Minneapolis, MN, USA]; NT3,[10 ng/ml, R&D Systems, Minneapolis, MN, USA]; db-cAMP [0.5 mM, Sigma-Aldrich, St. Louis, MO, USA]). ND medium was exchanged every 2-3 days. Small molecules and doxycycline were withdrawn from the medium after two weeks, to promote maturation of the neurons. The converted spheroids were kept in ND medium supplemented with growth factors, partially exchanged every 2-3 days, until the termination of the experiment. For 2D conversion, 24-well plates (Costar, Corning, USA) were serially coated with polyornithine (15 ug/ml, Sigma-Aldrich, St. Louis, MO, USA), laminin (5 ug/ml, Gibco, Thermo Fisher, Waltham, MA, USA) and Fibronectin (0.5 ng/ul, Gibco, Thermo Fisher, Waltham, MA, USA). 100.000 single hGPCs were seeded and after 24h they were transduced at MOI 1-2. After 24h, 500 μl of GM supplemented with doxycycline (2 μg/ml, Duchefa) were exchanged for activation of tet-regulated transgenes. Same media was used as in 3D cultures.

### Fluorescence-activated cell sorting (FACS) analysis

Two to three weeks after thawing, hGPCs were analyzed using FACS. Briefly, cells were mechanically detached and dissociated with Accutase (StemPro, Thermo Fisher; Waltham, MA, USA) for 8 min and were resuspended in Miltenyi wash buffer (PBS, Gibco, Thermo Fisher; Waltham, MA, USA; 0.5% BSA Fraction V Gibco, Thermo Fisher; Waltham, MA, USA; 2μM EDTA; 0.05‰ Phenol Red Sigma-Aldrich, St. Louis, MO, USA) in a concentration of 1×10^6^ cells/ml. This was followed by incubation with fluorochrome-conjugated antibodies (PE anti-human CD140a, [BD Biosciences, cat. no. 556002, Eysins, Switzerland 1:10]; APC anti-CD44, [Miltenyi, cat. no. 130-113-331, Bergisch Gladbach, Germany, 1:100]; APC anti-human CD133/1[Miltenyi, cat. no. 130-113-668, Bergisch Gladbach, Germany, 1:50]; FITC anti-human SSEA-4, [Biolegends, cat. no. 330410, Bergisch Gladbach, Germany, 1:20]) for 15 min at 4°C and washed in wash buffer for 10 min at 200 x g. Subsequently, cells were transferred into pre-wet 5 ml polystyrene tubes with cell-strainer caps with propidium iodide (PI, Miltenyi, cat. no. 130-095-177, 1:500, Bergisch Gladbach, Germany) to exclude dead cells. 10.000 events were analyzed on a FACSAria III sorter (BD Biosciences, Eysins, Switzerland). Gates were set on Fluorescence Minus One (FMO) controls and compensation was performed with single-stained cells.

### RNA extraction, cDNA synthesis, RT-qPCR

Spheroids were lysed with RLT buffer (Qiagen, Hilden, Germany), and total RNA extraction was performed using RNeasy Micro Kit (Qiagen, Hilden, Germany)) following the manufacturer’s protocol. Subsequently, RNA was reverse transcribed to cDNA using Maxima First Strand cDNA Synthesis Kit (Thermo Fisher, Waltham, MA, USA) according to the manufacturer’s protocol. The RT-qPCR mix was pre-mixed using the Bravo Automated Liquid Handling Platform (Agilent, Santa Clara, CA, USA). cDNA (1 μl) was mixed with LightCycler 480 SYBR Green I Master (5 μl, Roche, Basel, Switzerland) and relevant primers (4 μl, Eurofins Genomics, Luxembourg City, Luxembourg; See table S3) in 3 technical replicates for each sample. RT-qPCR was performed in a LightCycler 480 II instrument (Roche, Basel, Switzerland) using a 40 cycles two-step PCR protocol (95°C, 30s denaturation and 60°C, 1 min annealing/elongation). Analysis was performed from technical triplicates and the relative gene expression was calculated using the comparative CT Method (ΔΔCT Method) and every sample was compared to hGPCs before lentiviral transduction. Expression was normalized against the housekeeping genes *GAPDH* and *ACTB*.

### Whole spheroid immunostaining with optical clearing and 2D immunostaining

Whole spheroid immunostaining and optical clearing were performed according to^35^. Briefly, converted and control spheroids were fixed using 4% paraformaldehyde (PFA) or a mixture of 4% PFA and 0.25% glutaraldehyde (cat. no. G6257, Sigma-Aldrich, St. Louis, MO, USA) for 20 min at room temperature (RT). An overnight incubation in blocking solution containing 5% donkey serum, 0.3% Triton X-100 (Sigma-Aldrich, St. Louis, MO, USA) and PBS followed. Incubation with primary antibodies diluted in blocking solution was performed for 24h at RT on a shaker (See Table S4). This was followed by incubation with secondary antibodies conjugated to Alexa-488/Cy2, Alexa-568/Cy3, or Alexa-647 (1:200; Jackson ImmunoResearch Laboratories, West Grove, PA, USA) and DAPI (1:2000, Sigma-Aldrich, St. Louis, MO, USA) for 24h at RT on a shaker. Samples were then dehydrated in an ascending serial methanol concentration for 10 min each and lipids were removed by incubation with a dichloromethane-methanol mixture for 1h, followed by 2 steps of 10 min incubation, each in dichloromethane. Finally, spheroids were cleared with ethyl cinnamate and transferred to 96 well-plates with flat and clear bottoms (Ibidi, Gräfelfing, Germany) for confocal microscopy.

For 2D immunostaining, cells were fixed with 4% PFA for 15 minutes in RT. Subsequently, cells were incubated in a blocking solution (5% donkey serum, 0.1% Triton-X 100 in PBS) for 1-3h. Cells were incubated with primary antibodies (Table S4) diluted in a blocking solution overnight at 4°C. The next day, incubation with secondary antibodies and DAPI was performed for 1h at RT in the dark. Cells were kept at 4°C until microscopy.

### Patch clamp electrophysiological recording of converted spheroids

Whole-cell patch-clamp electrophysiological recordings were performed at 7-, 13-, 21- and 50-days post-transduction. Briefly, converted free-floating spheroids were recorded at RT in Krebs solution containing (in mM): 119 NaCl, 2.5 KCl, 1.3 MgSO_4_, 2.5 CaCl_2_, 25 Glucose and 26 NaHCO_3_ continuously gassed with 95% O2–5% CO_2_. Borosilicate glass pipettes (5–7 MOhm) filled with the following intracellular solution (in mM): 122.5 K-gluconate, 12.5 KCl, 0.2 EGTA, 10 Hepes, 2 MgATP, 0.3 Na_3_GTP, and 8 NaCl adjusted to pH 7.3 with KOH were used. Recordings were carried out with Multiclamp 700B System (Molecular Devices), digitized at 20 kHz, and acquired with pClamp 10.6 (Molecular Devices). Pipette access resistance was monitored throughout the recording and was between 9 - 25 MΩ. Cells were selected based on viability, neuronal shape, and clean access. Immediately after whole-cell configuration was established, resting membrane potential (RMP), input resistance (Ri), and cell membrane capacitance (Cm) were calculated from a series of 5 mV pulses of 100 ms duration. Action potential generation was examined by delivering a series of 500 ms duration depolarizing steps starting from −20pA to +35 pA with 5pA increments at current clamp while the RMP of the cells were held at –70 mV by constant current injection. Peak amplitude and afterhyperpolarization of the first elicited action potential were calculated as the voltage difference from the action potential threshold. For inward sodium (Na+) and delayed rectifying potassium (K+) current measurements, cells were clamped at −70 mV, and 100ms duration depolarizing steps were delivered starting from -70mV to +40mV with 10 mV increments at voltage-clamp. Igor Pro 8.04 (Wavemetrics), combined with the NeuroMatic package, was used for data analysis.

### Optical photothermal infrared microspectroscopy (O-PTIR)

O-PTIR experiments were performed using the mIRage^TM^ microscope (Photothermal Corp., St. Barbara US) available at SMIS beamline at the synchrotron SOLEIL(France). For these measurements, 20μm thick cryo-cut spheroids were directly deposited on a glass slide. The photothermal effect was detected by modulating the intensity of a continuous wave (CW) 532 nm laser, induced by an infrared laser. The probe power was set to 2.4%. Hyperspectral images were acquired by scanning from 3026 to 1000 cm^−1^ at an 80-kHz repetition rate using 2μm x 2μm step. Further details on the spectroscopic measurements of tissue can be found in previous work^36^. Data analysis was performed using Quasar^TM^ software using for unsupervised analysis of spectra^37^. The preprocessing of spectra involved cutting out the spectral part from 1300 to 1000 cm^−1^ to avoid glass sample support contribution; followed by baseline correction and min-max normalization. Spectra of poor quality (with high signal-to-noise ratio) were excluded using k-means clustering. Subsequently, principal component analysis (PCA) was used to investigate characteristic signatures. Specifically, we focused on the lipid signature within the 3026–2800 cm^−1^ range, the protein signature within the 1700– 1500 cm^−1^ range, and metabolites within the 1475–1300 cm^−1^ range.

### Nuclei Isolation and sorting

Spheroids were collected at different time points during the experiment, snap-frozen on dry ice and kept at -80°C until processed. Nuclei isolation was performed according to the following protocol^38^ with modifications. Briefly, spheroids were thawed at 4°C and dissociated in ice-cold lysis buffer consisting of 0.32 M sucrose, 5 mM CaCl_2_, 3 mM MgAc, 0.1 mM Na_2_EDTA, 10 mM Tris-HCl pH 8.0, 1 mM DTT, 0.1% Triton X, EDTA-free proteinase inhibitor (Roche, Basel Switzerland) and RNAse inhibitors (Ambion TM and SUPERase InTM, Invitrogen, Carlsbad, CA, USA). This was followed by centrifugation of the lysates for 30 min at 11.000 x g. The pelleted nuclei were resuspended in a buffer containing 0.1% BSA Fraction V, PBS -/-, RNAse inhibitors (Ambion TM and SUPERase InTM, Invitrogen, Carlsbad, CA, USA) and Draq7™ (BD Biosciences no. 564904, Eysins, Switzerland). The nuclei were filtered with a 70 μm cell strainer into BSA-coated DNA LoBind tubes (Eppendorf, Hamburg, Germany) for sorting. FACS was performed with a FACSAria cell sorter on 100 μm nozzle and with FACSDiva software (BD Biosciences, Eysins, Switzerland) on a low flow rate to separate single nuclei. 10.000 nuclei were collected in each sample to a total volume of 20 μl and directly processed for cDNA libraries’ generation.

### snRNA-seq library preparation, sequencing and raw data processing

Single nuclei suspensions were loaded onto 10x Genomics Single Cell 3′ Chips (v 3.1) with the master mix according to the manufacturer’s protocol (https://support.10xgenomics.com/single-cell-gene-expression/index/doc/technical-note-chromium-single-cell-3-v3-reagent-workflow-and-software-updates) for the Chromium Single Cell 3′ Library to generate single nuclei gel beads in emulsion (GEMs, v3 chemistry). The libraries were sequenced on a NovaSeq 6000 according to this programme: 28 cycles of Read1, 98 cycles of Read2, and 8 cycles of Index1 with a 200-cycle kit. Raw base calls were demultiplexed and converted to fastq files with cellranger mkfastq program (bcl2fastq 2.20/cellranger 6.0) for subsequent processing and analysis.

### Bioinformatic analysis

The raw snRNA data was mapped using 10X cellranger with the reference transcriptome GRCh38-2020-A. Scanpy^39^ (version 1.9.3 with Python version 3.10.12) was used for the downstream analysis of the snRNA data. Cells with fewer than 1000 genes or more than 5% mitochondrial count were filtered out. Doublet detection and removal was performed using Scrublet^40^. After filtering, median UMI count was 3516 and the number of detected genes was 33512. Log-transformation was performed, and 3642 highly variable genes were identified. The Scanpy implementation of PCA dimensionality reduction was then used on the highly variable genes plus a selection of genes relevant to the desired conversion. For UMAP projection and clustering, the 20 nearest neighbors were used and the minimum distance set to 0.05. For integration with the published dataset, the Scanpy implementation of Harmony was used, and converged after 7 iterations. Differential gene expression analysis was done using Wilcoxon rank sum test as implemented by Scanpy, and genes with an FDR-adjusted p-value < 0.05 were considered significant. The python package Gseapy^41^ was used for gene set enrichment analysis on the top 25 differentially expressed genes, with the following databases: ”GO Biological Process 2023”, ”GO Cellular Component 2023”, and ”GO Molecular Function 2023”, as well as the corresponding versions from 2021. For RNA velocity analysis, the Python package Scvelo^42^ was used. Plots were produced with the Scanpy, Scvelo, Gseapy, and Matplotlib Python packages.

### Microscopy and PV quantification

Fluorescent images were taken with a Zeiss 780 Confocal Laser-Scanning Inverted Microscope (20X objective and 40x object) and with Zeiss Zen Blue Edition software. For the 40X pictures, the HDR mode was used. Images were edited on ImageJ (NIH, Bethesda, MA, USA) and the changes were applied equally across the entire image without loss of information. Quantification of PV expressing cells in spheroids was performed in ImageJ (NIH, Bethesda, MA, USA) by manually counting the individual PV+ cells in all the 1µm-thick z-stacks of the confocal images. Brightfield images were captured with an Olympus IX50 Inverted Phase Contrast Microscope and acquired with CellSens Standard software.

### High-content screening

The neuronal purity in 2D was manually calculated as the number of TAU+ cells over the DAPI+ nuclei using Operetta High Content Imaging System and the Harmony 5.2 software (Waltham, MA, USA). For each experiment, 371-379 fields (20X magnification) from one well were analyzed. The measurement of the mean fluorescence for antigen-positive cells was defined above the background fluorescence of antigen-negative cells (internal negative control). The same strategy was used for the analysis of the number of PV+ cells containing DAPI+ nuclei and expressing TAU. Using the “Neurite Outgrowth” script on Harmony 5.2 (Perkin Elmer, USA), the total neurite length and segment length were quantified for PV+/TAU+ neurons.

### Statistical analysis

Statistical analyses were performed using GraphPad Prism 9.4.1. A Shapiro-Wilk normality test was applied to assess the normality of the distribution, and parametric and non-parametric tests were performed accordingly. PV+ cells quantifications, RT-qPCR and electrophysiology data were compared using either one-way ANOVA followed by post-hoc Tukey test or non-parametric Kruskal-Wallis test and uncorrected Dunn’s test. RT-qPCR data, neurite analysis and PV quantification on Figure S6 were compared using either unpaired t-test or Mann-Whitney test. All data are expressed as mean ± standard error of the mean (SEM). The level of statistical significance was set at p<0.05.

## Results

### Neuronal reprogramming of human glia precursor cells in 3D spheroids

hGPCs were cultured for 2-3 weeks after thawing and then seeded in ultra-low attachment plates along with lentiviral vectors carrying five reprogramming factors (*Ascl1*, *DLX5*, *LHX6*, *Sox2* and *FOXG1*; ADLSF) in a dual regulation system^30^ (Figure 1A). A human embryonic stem cell (hESC) differentiation protocol was used for generating hGPCs^28^ according to the outline in Figure 1A. Before reprogramming, hGPCs exhibited typical bipolar morphology (Figure 1B) and expressed Platelet Derived Growth Factor Alpha (PDGFRα), marking migrating oligodendrocyte precursor cells, as well as Glial Fibrillary Acidic protein (GFAP), labelling astrocyte biased precursors or bipotent precursors, when co-expressed with PDGFRα (Figure 1C). After seeding, cells rapidly self-aggregated into 3D spheroids and were cultured for up to 50 days (Figure 1A, D). For control groups, hGPCs were kept without reprogramming factors in either glial medium (GM) or neuronal differentiation medium (ND) for 21 days (Figure 1A).

**Figure 1.**
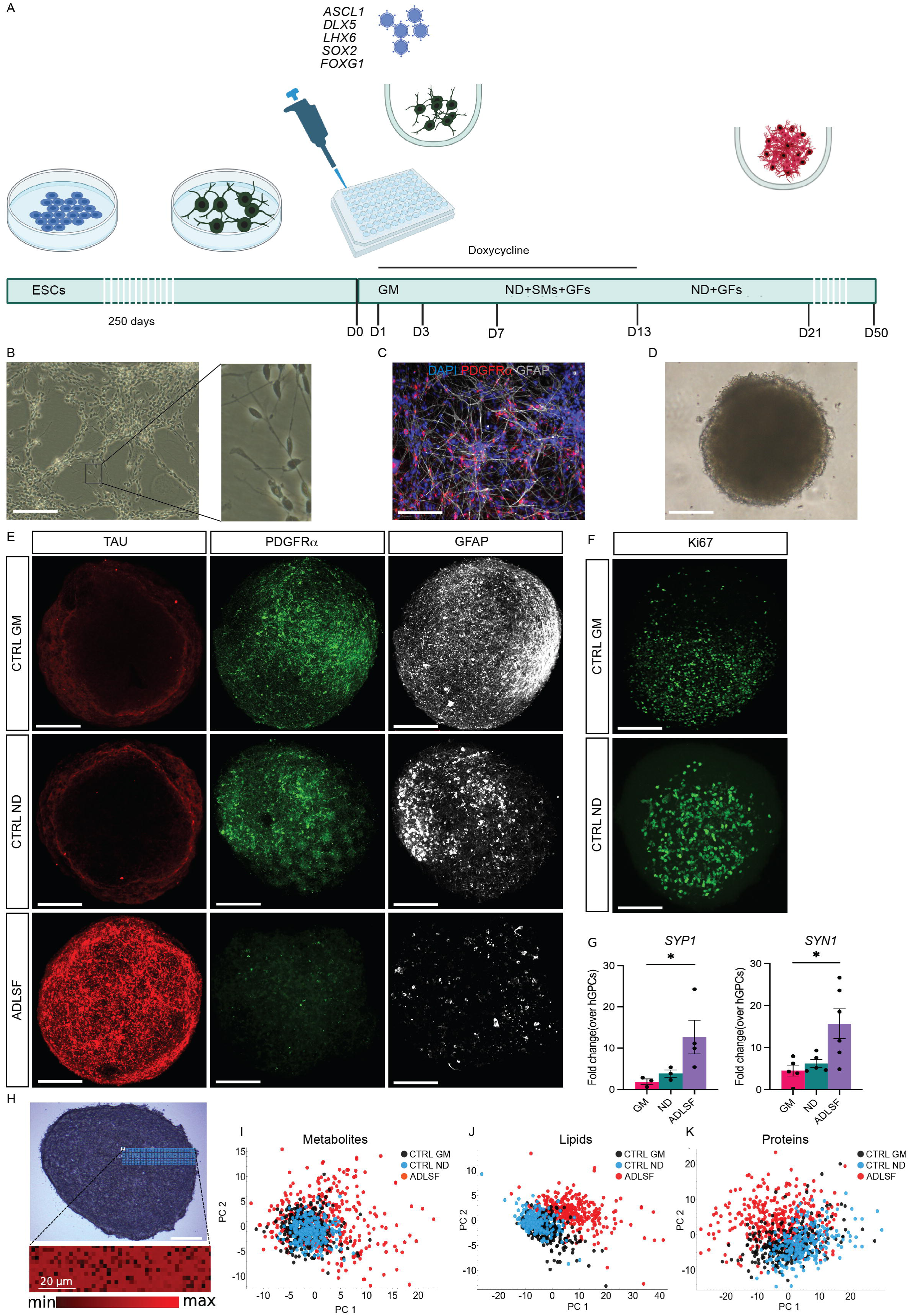
Neuronal reprogramming of human glia precursor cells in 3D spheroids. A) Schematic overview of the programming process. B) A brightfield photo of hGPCs in 2D culture at day 280 of differentiation. C) Immunocytochemistry of hGPCs demonstrates the presence of glia markers PDFGRα and GFAP. D) A brightfield photo of an induced 3D spheroid. E) Maximum intensity projections of immunostainings for TAU, PDGFRα and GFAP in ADLSF-transduced cells and glial cells in GM (glial medium) and ND (neural differentiation medium). F) Maximum intensity projections of immunostaining with the proliferation marker MKi67 in GM and ND controls. G) RT-qPCR analysis of *SYN1* and *SYP* (n=3-5 for GM and ND, n=3-6 for ADLSF n=biological replicate): *p < 0.05. One-way ANOVA test and post-hoc Tukey test was performed for SYN1 and Kruskal-Wallis with uncorrected Dunns’s test for *SYP1.* H) A bright field overview of induced spheroid. Blue dots indicating spectral positions and intensity distribution at 1656 cm⁻¹. Black spectra corresponding to low intensity were removed. I) PCA analysis of different biological replicates of the ADLSF condition. J) PCA analysis of lipids and proteins in the ADLSF condition compared with ND and GM control groups. Scale bar: 100um. Data are presented as mean ± SEM.

An initial immunocytochemistry assessment of the spheroids on day 21 demonstrated a clear induction of the neuronal marker TAU (microtubule-associated protein) in the ADLSF group that was not seen in GM or ND (Figure 1E). Along with this, there was a complete decline of PDGFRα protein and GFAP in the ADLSF group, but not in the GM and ND control, indicating a neuronal switch in the ADLSF group and remaining glial identity in controls (Figure 1E). The hGPCs in the two control groups remained proliferative, as shown by the Ki67 staining, further confirming their glial identity (Figure 1F). In line with the neuronal protein induction in ADLSF, RT-qPCR assessment showed upregulation of the pan-neuronal markers Synaptophysin 1 (*SYP1*) and Synapsin 1 (*SYN1*) compared to controls (Figure 1G). Furthermore, we could confirm by RT-qPCR the presence of all the individual transgenes at day 7,13 and 21 in the ADLSF transduced group (Figure S1A), highlighting the contribution of the viral cocktail to the neuronal conversion.

To confirm the phenotype switch in the 3D spheroid, we exclusively applied optical photothermal infrared (O-PTIR) microspectroscopy on ADLSF and controls from spheroid cryosections (Figure 1H). Spectra were taken from both outer and inner zones of individual spheroids with similar cell viability indicating homogeneity throughout the spheroid and good cell viability also in the core (Figure 1H). Importantly, analysing spectral intensities, we could detect a change in intracellular biomolecules such as metabolites, lipids and proteins upon reprogramming, i.e. in ADLSF condition compared to controls (Figure 1I, J and K), indicating a compositional cell change upon neuronal reprogramming.

Altogether, these data demonstrate successful neuronal conversion in 3D culture system that is transcription factor dependent and changes the biomolecular composition in the cells.

### Glia reprogramming show early fate dynamics with gradual functional neuronal state

To unravel the transcriptional dynamics of hGPC-to-neuron reprogramming we performed 10X Genomic droplet-based single nuclear RNA sequencing of 3D reprogrammed cells at 0,1,3,7,13 and 21 days post conversion. In total, 21,276 single nuclei were profiled after quality filtering from one biological replicate (Table S2). At day 0, before virus administration, the cells expressed classical glial markers *PDGFRA*, Protein Tyrosine Phosphatase Receptor Type Z1 (*PTPRZ1*), SRY-Box Transcription Factor 9 (*SOX9*), and astrocyte biased markers (*GFAP*) as seen on a uniform manifold approximation and projection (UMAP) analysis (Figure S2A), confirming a glial starting cell population ^43^.

From this state onwards, several transcriptionally distinct clusters could be identified throughout the reprogramming process until day 21 (Figure 2A and B). Distinct gene dynamics occurred as early as on day 3 and 7. New transcriptionally distinct clusters gradually appeared with a full transcriptomic switch at day 7 to 13. On day 21, another cluster took shape, which seemed transcriptionally closer to the earlier time points, probably attributed to the decreased cell yield for this time point due to the nuclei extraction procedure (Table S2). The neuronal reprogramming process did not seem to involve much apoptosis as low levels or no expression was detected for Caspase 3 (*CASP3*), BCL2 Associated X (*BAX*) and Fas Cell Surface Death Receptor (*FAS*) (Figure S2B). Furthermore, very few cells expressed cycling genes (*TOP2A*) (Figure S2B).

**Figure 2.**
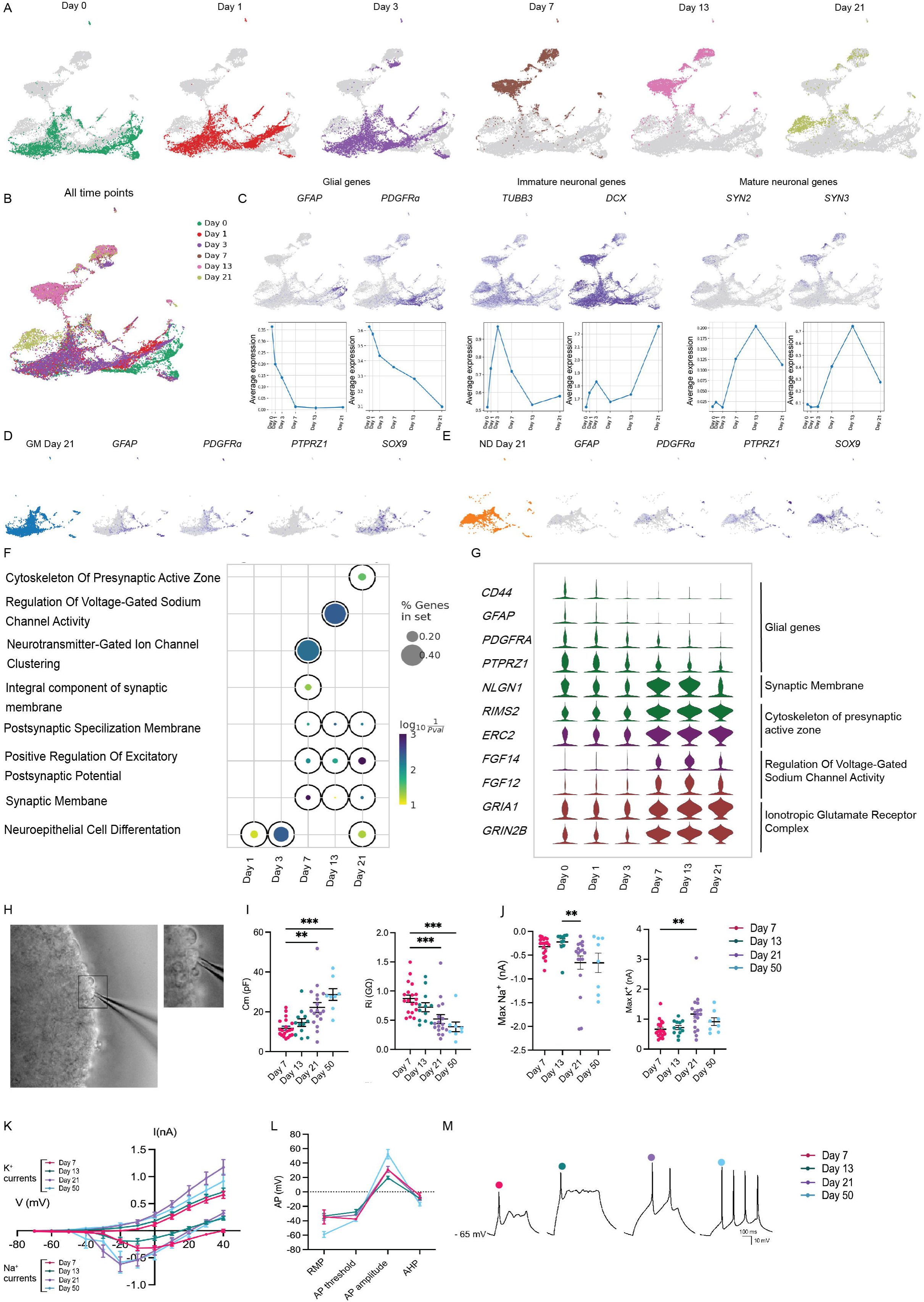
Glia reprogramming show early fate dynamics with gradual functional neuronal state. A) UMAP plots at different timepoints (day 0,1,3,7,13 and 21) during the reprogramming process. B) UMAP plot with all the time points together. C) UMAP plots of glial (*GFAP, PDGFRA*), immature neuronal (*TUBB3 and DCX*) and mature neuronal (*SYN2, SYN3*) genes with average expression levels over time (See also figure S2B, C). D, E) UMAP plots visualizing the expression levels of glial genes *GFAP, PDGFRA, PTPRZ1, SOX9* for D) Glial medium (GM) control at day 21 and E) Neural Differentiation control (ND) at day 21 (See also figure S2D). F) Pathway analysis of induced spheroids at different time points. G) Violin plot showing differentially expressed genes at day 0 and various time points during conversion. H) A brightfield photo of patch-clamp electrophysiology of a free-floating induced spheroid. I) Passive membrane properties of induced spheroids at different time points (n=19-20 for day 7, n=12 for day 13, n=17-18 for day 21, n=8 for d50, n=individual cell): * p < 0.05, **p<0.01, ***p<0.005. Kruskal-Wallis test with uncorrected Dunns’s test. J) Maximum sodium (Na+) and maximum potassium (K+) currents at different time points demonstrating an increase in current over time (n=20 for day 7, n=12 for day 13, n=18 for day 21, n=8 for day 50, n=individual cell): **p<0.01, Kruskal-Wallis test with uncorrected Dunns’s test. K) Inward sodium (Na+) and outward potassium (K+) currents plotted against stepwise voltage induction (right), maximum Na^+^ current (middle) and maximum K^+^ current (left) (n=20 for day 7, n=12 for day 13, n=18 for day 21, n=8 for day 50, n=individual cell). L) Action potential (AP) properties, resting membrane potential (RMP), AP threshold, AP amplitude and after-hyperpolarization (AHP) (n=2 for day 7, n=3 for day 13, n=11 for day 21, n=3 for day 50, n=individual cell). M) Representative traces at different time points. Each dot represents the mean value. Electrophysiology recordings were obtained from 3 biological replicates. Data are presented as mean ± SEM.

The dynamic transcriptional change followed a glia-to-neuron transition as seen with specific marker genes of glia cells, immature neurons and mature neurons (Figure 2C). Glial genes *GFAP* and *PDGFRA,* were upregulated in the earlier time points and gradually downregulated during the process, whereas the immature neuronal genes Tubulin Beta 3 Class III (*TUBB3*) and doublecortin (*DCX*) were transiently expressed at day 3 and then declining by day 7 (Figure 2C). Only DCX re-emerged at day 21, potentially at the expense of reduced mature neuronal population at this time point (Table S2)). The mature neuronal genes Synapsin II and III (*SYN2*, *SYN3*) were mainly detected in the clusters occurring from day 7 and day 13 (Figure 2C).

During the reprogramming process, there was concurrent expression of endogenous transcription factors (*DLX5, SOX2,* and *FOXG1*), mirroring those used in the viral transduction most likely due to a reinforcing action of transgenes (Figure S2C).

In contrast to the experimental condition, the control hGPCs maintained their glial state, as marked by the expression of e.g. *GFAP* (3% in GM and 1.4% in ND), *PDGFRA* (9.9% in GM and 6% in ND)*, SOX9* (11.1% in GM and 13.2% in ND), and *PTPRZ1* (23.6% of GM and 21.7% in ND) (Figure 2D, E). A slightly different cell population was seen in ND control, compared to GM control (Figure 2E). This was possibly an effect of long-term ND medium with small molecules and growth factors, influencing the fate of these cells (Figure 1A). Nevertheless, this population continued to express glial markers, further highlighting the necessity of transcription factors for neuronal conversion. Furthermore, few cells expressed the cycling gene *TOP2A* in day 0 and both controls. (Figure S2D). Altogether, this demonstrates a distinct glia-to-neuron transcriptional process in 3D reprogrammed glia and the necessity of ADLSF transduction for successful neuronal conversion.

Pathway analysis on the reprogrammed cells from day 1 to 21 confirmed the acquisition of a neuronal fate, switching from immature pathways, e.g. Neuroepithelial cell differentiation pathway into more neuronal specific pathways from day 7, e.g. Neurotransmitter-Gated Ion Channel, Voltage-Gated Sodium Channel activity, and Excitatory Postsynaptic Potential regulation (Figure 2F). Interestingly, the Neuroepithelial cell differentiation pathway reappeared at day 21, further supporting the previous data of neuronal loss at this time point. This data supports a rapid glia conversion to a mature and functional neuronal fate.

Next, we sought to investigate the reprogrammed cells for differentially expressed genes involved in these pathways. Here, we could further confirm a glial fate at earlier time points (day 0-3), with expressions of *CD44*, *GFAP*, *PDGFRA* and *PTPRZ1* (Figure 2G). From day 7 to 21 instead, genes involved in pivotal neuronal processes appeared, e.g. synaptic membrane compartment (Neuroligin 1: *NLGN1*) (Figure 2G). Presynaptic active zone genes, such as Regulating Synaptic Membrane Exocytosis 2 *(RIMS2*) and synaptic membrane genes ELKS/RAB6-Interacting/CAST Family Member 2 (*ERC2)* were strongly induced from day 7 along with a functional neuronal state and synapse formation (Figure 2G). Genes related to Voltage-Gated Sodium Channel Activity, such as Fibroblast Growth Factor 14 *(FGF14*) and Fibroblast Growth Factor 12 *(FGF12*), were upregulated from day 7, with highest expression at day 13 whereas genes involved in excitatory postsynaptic potential e.g. AMPA or NMDA receptors (*GRIA1, GRIN2B*, encoding for GluN2B subunit), were more highly expressed from day 7 to 21 (Figure 2G), supporting the formation of synaptic compartments and functional neuronal membrane properties. Altogether, this pathway analysis demonstrates a transcriptomic switch from a glia precursor state to a functional neuronal state with synaptic compartment formation in only 13 days of reprogramming.

To further prove a functional neuronal state, we performed patch-clamp electrophysiology on free-floating spheroids from day 7 up to 50 days of culture (Figure 2H and Table S4). This showed a gradual functional improvement of the intrinsic membrane properties throughout the reprogramming process. The input resistance (Ri) was reduced at day 13 and 21 compared to day 7, indicating an increased number of voltage gated channels and membrane capacitance (Cm) was significantly increased from day 21, indicating increased cell size (Figure 2I). At all time points, rectifying inward sodium (Na^+^) and outward potassium (K^+^) were detected (Figure 2J) with a gradual increase in Na^+^ amplitude at day 21 and day 50 (Figure 2K). The resting membrane potential (RMP) was more hyperpolarized at day 50 compared to day 13. Whereas action potential (AP) threshold was similar among time points, the AP amplitude and afterhyperpolarization (AHP) differed between day 50 and the earlier time points, suggesting a gradual functional maturation in firing properties (Figure 2L). Furthermore, the ability to fire repetitive series of evoked APs upon current injection was superior at day 50 compared to that seen at day 21 (Figure 2M). Overall, this data indicates a gradually maturation in neuronal function in the reprogramming process, in line with the protracted maturation process of neurons *in vitro*^10^.

### hGPCs convert into GABAergic interneurons with parvalbumin phenotype

Based on the sequencing analysis above, we decided to focus the analysis on the neuronal clusters with the appearance of three transcriptionally distinct neuronal clusters labelled in yellow, purple and green (Figure 3A). To further understand the cell phenotypes in these neuronal clusters, we performed an unbiased analysis for the highly expressed genes (Figure S3A). This analysis revealed that the most highly expressed gene in green and purple clusters was the long non-coding RNA *SOX2-OT,* mainly expressed in GABAergic interneurons^44^ and Small Nucleolar RNA Host Gene 14 (*SNHG14,)* respectively, mainly edited on RNA level, in interneurons^45,46^. Other highly expressed genes included various cell adhesion molecules; Neurexin 1 and 3 (*NRXN1, NRXN3*) in green cluster, Neurotrimin (NTM) and Cell adhesion Molecule 1 (*CADM1*) in purple cluster and Limbic System Associated Membrane Protein (*LSAMP*) and FERM Domain Containing 5 (*FRMD5*) in yellow cluster (Figure A3A).

**Figure 3.**
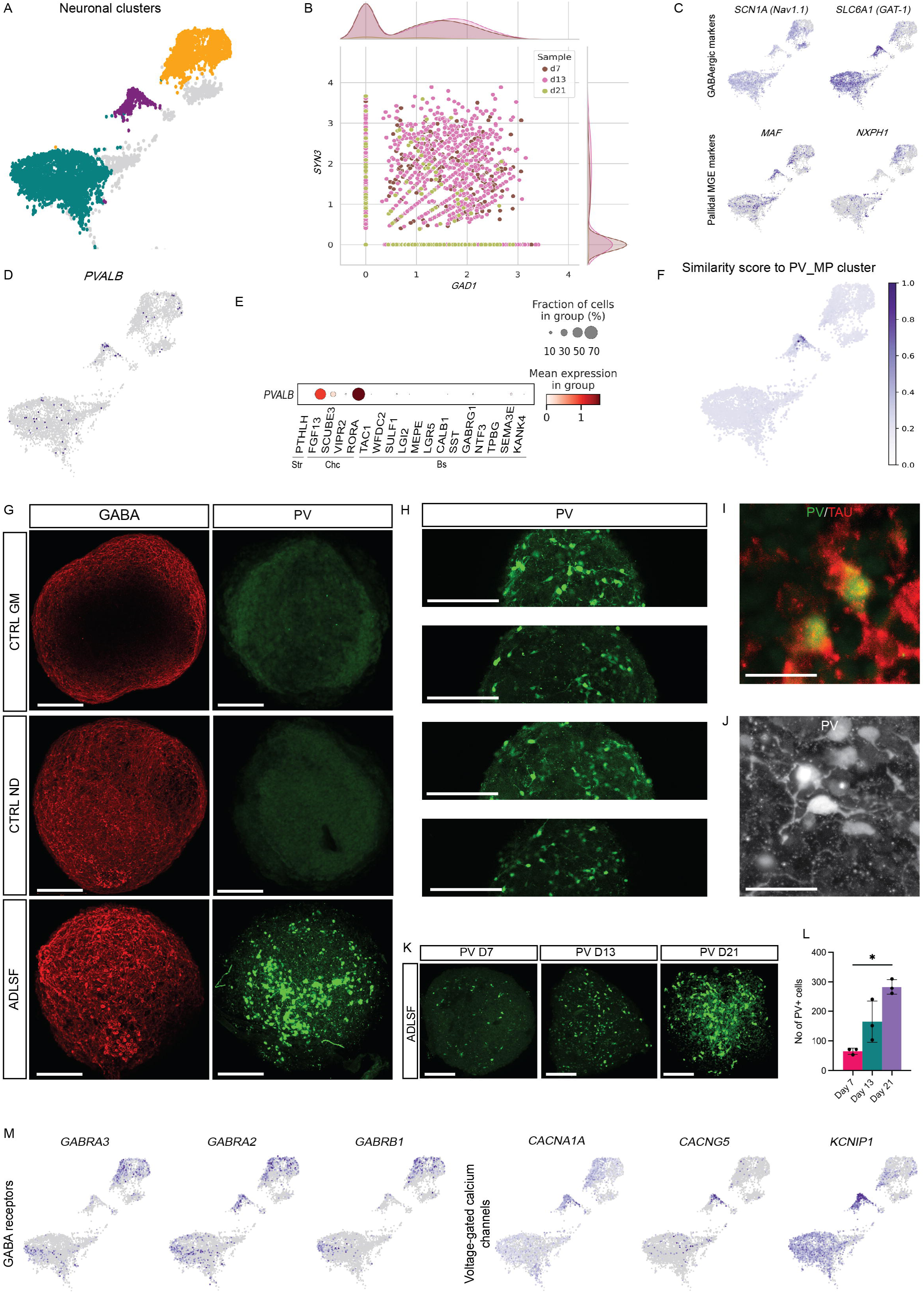
Induced neurons are primarily GABAergic interneurons with PV phenotype. A) UMAP plot visualizing the neuronal clusters (See also figure S3A). B) Scatter plot of co-expression of *GAD1* and *SYN3* in the neuronal clusters. C) UMAP plots demonstrating the expression levels of pan-GABAergic markers and pallidal GABAergic markers (See also figure S3B). D) UMAP plot of expression levels of *PVALB* (See also figure S3C). E) Dot plot showing co-expression of *PVALB* with subclass-specific markers. F) UMAP plot of cosine similarity score between the PV_MP cluster of the published dataset^53^ and the PV cluster in this dataset (See also figure S3D). G) Maximum intensity projections of immunostaining of GM and ND controls and induced spheroids with GABA and PV (See also figure S4A). H) Different z-stacks of immunostaining of PV and its distribution throughout the whole spheroid. I) Immunostaining of PV and pan-neuronal marker TAU in 3D spheroids. J) 40X picture confocal image of a single PV interneuron. K) Representative images of immunostaining of PV cells at day 7,13 and 21. L) Quantification of PV cells at day 7,13 and 21 (n=3, n=biological replicate): * p<0.05, Kruskal-Wallis test with uncorrected Dunns’s test. M) UMAP plots with expression levels of GABA receptors (*GABRA3*, *GABRA2* and *GABRB1*) and voltage-gated calcium channels (*CACNA1A, CACNG5* and *KCNIP1*) Scale bars: in G, H, K): 100μm. Scale bar in I, J) 40μm. Each dot represents a biological replicate. Data presented as mean ± SEM.

We further evaluated the three neuronal groups for the expression of Glutamate Decarboxylase 1 (*GAD1*), the enzyme responsible for GABA production, and its co-expression with the mature neuronal marker *SYN3* (Figure 3B). A scatterplot revealed that approximately two-thirds of *SYN3* cells co-expressed *GAD1* at day 7 to 21 (Figure 3B, n=1274 cells out of a total of 1888 *SYN3*-expressing cells), indicating an inhibitory neuronal population. Fewer cells expressed glutamatergic marker, e.g., Solute Carrier Family 17 Member 7 (*SLC17A7), VGLUT-1,* Protein Phosphatase 1 Regulatory Inhibitor Subunit 1B (*PPP1R1B) and* medium spiny neuron marker DARPP-32 (Figure S3B).

Further analysis demonstrated expression of selective GABAergic markers e.g. Sodium Voltage-Gated Channel Alpha Subunit 1 Nav1.1 (*SCN1A*), mainly expressed in GABAergic interneurons^47^, and Solute Carrier Family 6 Member 1 (*SLC6A1*) or *GAT-1,* mainly expressed in PV chandelier interneurons^48^ (Figure 3C). Moreover, we could confirm an MGE-derived pallial fate, as shown by the expression of MAF BZIP Transcription Factor (*MAF*) and Neurexophilin 1 (*NXPH1*)^9^(Figure 3C). We could further exclude the presence of other MGE-derived fates, investigating LIM Homeobox 8 (*LHX8*) and Choline O-Acetyltransferase (*CHAT*), markers of cholinergic interneuron identity, that showed expression in very few cells, possibly reflecting a small off-target effect from the transcription factors (Figure S3B).

Analysis of interneuron subtype specific markers interestingly showed that parvalbumin (*PVALB*) was the most prominent subtype specific marker expressed among the clusters and specifically enriched in purple cluster (Figure 3D). Other subtype-specific markers (e.g. *SST, CALB1, VIP and CALB2*) were expressed in fewer cells compared to PV cells (Figure S3C). We further explored three distinct categories; striatal, cortical chandelier and cortical basket PV interneurons, each characterized by specific marker expression and connectivity^48–52^. Results showed that most PV cells belonged to the chandelier subclass, with 50% of PV cells co-expressing Fibroblast Growth Factor 13 (*FGF13),* 30% co-expressing Signal Peptide, CUB Domain And EGF Like Domain Containing 3 (*SCUBE3*), and 70% co-expressing RAR Related Orphan Receptor A (*RORA*). No PV cells co-expressed the striatal marker Parathyroid Hormone Like Hormone (*PTHLH*), which marks striatal PV cells, and a few PV cells expressed markers for basket cells such as Extracellular Sulfatase Sulf-1 (*SULF1*), Semaphorin 3E (*SEMAE3*) and Gamma-Aminobutyric Acid Type A Receptor Subunit Gamma1 (*GABRG1*) (Figure 3E).

To test if the induced PV interneurons were equivalent to *bona fide* human endogenous interneurons, we compared our dataset with a recently published dataset of cortical fetal and adult human interneurons^53^. Using the top 50 principal components, we compared the induced PV neurons with the endogenous PV_MP cluster i.e. the PV chandelier branch of the published dataset^53^. This comparison produced a cosine similarity score of 0.8 out of maximum of 1, highlighting the high similarity between our cells to the endogenous PV subgroup when compared with the rest of the clusters from the dataset (Figure 3F, S3D). Interestingly, the four clusters with the highest similarity scores to our PV cluster were all MGE-derived interneurons, further supporting the presence of MGE-fate cells (Figure S3D). These results show for the first time that human induced PV interneurons can have a transcriptional subtype and subclass characteristics equivalent to endogenous adult PV interneurons.

To confirm the sequencing results, we assessed the protein expression of GABA and PV using immunocytochemistry (Figure 3G). While GABA protein expression was low in the ND and GM controls, it was drastically increased in the ADLSF condition at day 21 (Figure 3G bottom panel). Moreover, there was a clear induction in PV protein levels in the ADLSF condition, which was not observed in the control groups, proving that the transcription factors are necessary for successful PV conversion.

We additionally tested reprogramming with fewer transcription factors i.e. ADL to reduce the viral load to the hGPCs and to unravel the need of all five factors but that did not produce similar PV reprogramming efficiency (Figure S4A,B), suggesting superior outcome with ADLSF combination.

The induced PV cells were detected throughout the entire 3D spheroid structure as assessed by analyzing z-stacks from confocal imaging, demonstrating a homogenous PV reprogramming within the 3D structure (Figure 3H). Importantly, PV cells showed co-localization with the mature neuronal marker TAU (Figure 3I) and manifested a complex neuronal phenotype with distinct dendritic trees within the spheroid structure (Figure 3J), implying a mature neuronal phenotype. To observe the potential changes in PV protein expression over time, we manually quantified the PV cells at day 7 to day 21 post transduction. Quantification showed a gradual increase over time, reaching 283 ± 25 PV cells per spheroid at day 21 (Figure 3K, L). In line with the sequencing analysis, RT-qPCR analysis showed a clear upregulation of the pan-GABAergic marker Solute Carrier Family 32 Member 1 (*SLC32A1* or *VGAT*) and the interneuron subtype specific marker *PVALB* (Figure S3E). While there was a small induction of *SST* and *CALB1* genes in the RT-qPCR (Figure S3E), the expression levels were lower compared to the *PVALB* gene and could not be detected on the protein level. Together, these data indicate a homogenous and efficient induction into PV interneurons in the 3D culture system.

Next, we investigated genes encoding functional characteristics of PV interneurons, such as the presence of voltage-gated calcium channels important for their fast-spiking firing^54^. In all neuronal clusters, and particularly on the PV-enriched cluster, we found genes for different GABA receptor subunits, facilitating communication with other inhibitory neuron e.g. Gamma-Aminobutyric Acid Type A Receptor Subunit Alpha 2 (*GABRA2*), Subunit Alpha (*GABRA3)* and Subunit Beta1 (*GABRB1)*^55^ (Figure 3M). Moreover, Calcium Voltage-Gated Channel Subunit Alpha1 A (*CACNA1A*), and Calcium Voltage-Gated Channel Auxiliary Subunit Gamma 5 (*CACNG5*), which encode for the Voltage-Dependent Calcium channel of P/Q Type and Gamma-5 Subunit respectively, were expressed, along with Potassium Voltage-Gated Channel Interacting Protein 1 (KCNIP1) (Figure 3M). In conclusion, these data suggest that the induced GABAergic interneurons express ion channels reminiscent of cortical PV interneurons.

### Constructing the lineage trajectory of reprogrammed parvalbumin interneurons

To further understand the glia-to-PV conversion, we examined the cellular transcriptomic trajectory using latent time reconstruction of the single nuclear RNA sequencing data. We based this on RNA velocity^42^ with Figure 2B as a backbone. Latent time was used to trace cell maturity from a “branching point” (red cells in Figure 4A) i.e. glia cluster to the PV cluster (dark blue cells in Figure 4A, Figure S5A, B) with RNA velocity and PAGA^56^ analysis and revealed likely transitions between assigned glial cells and PV cells (Figure S4C). This analysis resulted in the generation of two distinct lineage trajectories. The left trajectory did not reach the PV cluster but instead led to other clusters, indicated as “non-PV trajectory” (black arrows in Figure 4A), while the right trajectory successfully targeted the PV cluster and was designated as the “PV trajectory” (blue arrows in Figure 4A).

**Figure 4.**
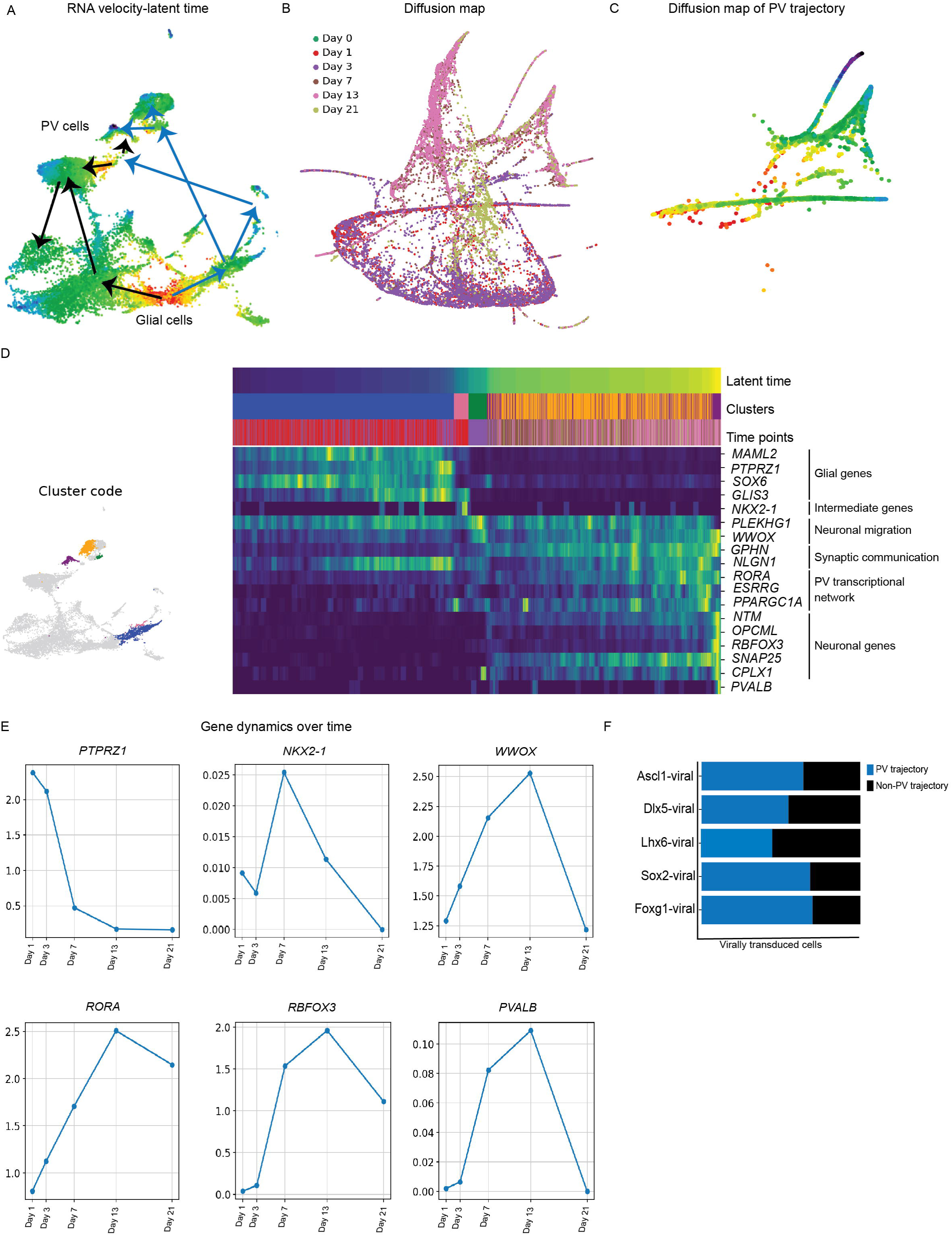
Constructing the lineage trajectory of human induced PV interneurons. A) UMAP visualizing the reconstructed reprogramming trajectory on latent time (See also figure S5A, B, C). B) Diffusion map with all time points (See also figure S5D). C) Representation of the reconstructed PV trajectory of the diffusion map colored by latent time. D) Clusters involved in the PV trajectory and heatmap visualizing important genes of the PV trajectory. E) Average expression of specific genes at different time points of reprogramming. F) Proportion of transduced cells belonging to the PV trajectory and non-PV trajectory respectively.

To further corroborate these findings from latent time-RNA velocity analysis, we performed diffusion pseudotime analysis^56^, which was based on changes in gene similarity (Figure 4B). This confirmed the presence of a distinct PV cluster (Figure S5C) and revealed a dedicated lineage pathway leading to the PV-enriched cluster. The cells within the PV trajectory appeared interconnected in the diffusion map, with the end of the PV-enriched cluster showing the highest level of maturity according to latent time, as indicated by the blue color (Figure 4C). Based on these findings we inferred that glia cell reprogramming diverges into two distinct lineages; one that does not reach full maturity and remain in a GABAergic immature state (black arrows in Figure 4A), and another that achieves full maturity and acquires the PV neuronal fate (blue arrow in Figure 4A).

Next, we explored the genes and pathways expressed along the “PV trajectory” and plotted them against latent time (Figure 4D). The heatmap demonstrated a gradual downregulation of genes associated with hGPCs’ identity, such as Mastermind Like Transcriptional Coactivator 2 (*MAML2), PTPRZ1,* SRY-Box Transcription Factor 6 (*SOX6)* and GLIS Family Zinc Finger 3 (*GLIS3*)^57^. Conversely, there was an increase in the expression of mature neuronal markers, such as *NTM*, Opioid Binding Protein/Cell Adhesion Molecule Like, (*OPCML*), RNA Binding Fox-1 Homolog 3 (*RBFOX3)* and Synaptosome Associated Protein 25 (*SNAP25)*, indicating the transition towards a neuronal fate. Notably, there was transient upregulation of progenitor genes, such as NK2 Homeobox 1 (*NKX2-1)*^58^ and neuronal migration genes, such as Pleckstrin Homology and RhoGEF Domain Containing G1 (*PLEKHG1*) and WW Domain Containing Oxidoreductase (*WWOX*) (Figure 4E,F).

During the conversion process, there was an upregulation of postsynaptic proteins, such as gephyrin (*GPHN*) and *NLGN1*. Towards the end of the latent time, there was an upregulation of PPARG Coactivator 1 Alpha (*PPARGCA1*), Estrogen Related Receptor Gamma (*ESRRG*), and *RORA.* Certain genes in the heatmap displayed distinct dynamics overtime in the heatmap, with some rapidly downregulating (*PTPRZ1*) or exhibiting more dynamic expression patterns (*NKX2-1*) (Figure 4E). Interestingly, there was a robust upregulation of *WWOX, RORA, RBFOX3 and PVALB,* as early as day 7, further corroborating the fast dynamics involved in glia-to-PV interneuron conversion (Figure 4E).

Notably, the expression of each individual transgene (ADLSF) was higher in the trajectory towards PV fate compared to the non-PV trajectories, as illustrated in (Figure 4F). A higher fraction of cells expressing all transcription factors was observed for all factors except *LHX6*. Particularly, 61% of cells transduced with *Ascl1* and 55% of cells transduced with *DLX5* were found in the PV trajectory. Furthermore, 44% of *LHX6*-transduced cells, 68% of *SOX2*-tranfected cells and 69% of *FOXG1* transduced cells were part of the PV-trajectory. This further supports the pivotal role of reprogramming factors in the PV fate acquisition. In summary, these results highlight a novel reprogramming process where cells progress through different developmental stages, obtaining the necessary machinery for synaptic communication and energy demand to ultimately attain a complete PV neuronal fate.

The unravelling of important transcription factors in the PV trajectory persuaded us to manipulate them *in vitro*, with the aim to increase the PV yield in the cultures. We chose *RORA,* a highly expressed gene at day 7, for this purpose. RORA transgene was delivered into a doxycycline-regulated lentiviral vector with a GFP reporter and was added to the reprogramming cocktail (ADLSF+R) for transduction in both 2D and 3D culture conditions (Figure S6A). An initial 2D experiment confirmed successful expression of the RORA-GFP in the hGPCs, with proof that cells incorporating the transgene also express PV (Figure S6B). Transduction of ADLSF+R did not impair the induction of complex PV neuron morphology, as seen in 2D cultures (Figure S6C) but appeared to induce slightly longer neurites and segments (Figure S6D). PV yield also showed a trend of increased yield in this condition (5.8%±1.2), compared to the ADLSF control (4%±1) (Figure S6E) indicating that the addition of RORA did not negatively affect the yield of PV cells, but intriguingly, possibly resulted in more complex PV cells and slightly higher PV yield.

RT-qPCR analysis in the 3D spheroid culture showed a trend of increased *SYN1* and *VGAT* expression in the ADLSF+R condition and confirmed a small increase in *PVALB* gene (Figure S6E). PV protein was seen at similar levels in both conditions (Figure S6F, G). Overall, these data suggest that ectopically delivered *RORA* has potential to increase neuronal complexity as well as the PV gene expression without affecting the overall number of PV cells.

## Discussion

PV interneurons precisely regulate local circuitry, brain networks and memory processing^2^ and their dysfunction associates to several neurological disorders characterized by network alteration and cognitive impairment, including Schizophrenia, Epilepsy, Alzheimer’s disease (AD), and autism spectrum disorders^3–7,59^

Experimental studies of interneuron transplantation have showed attenuation of disease pathology for AD, epilepsy or schizophrenia^5,7^ and potential for interneurons to reopen the critical window of cortical plasticity^60^. However, generating PV interneurons for cell transplantation, whether from stem cells or aborted fetuses, has proven to be difficult or even unfeasible due to limitations in cell sourcing^13^.

Herein, we have established a new approach to generate human PV interneurons by reprogramming hGPCs that shows promise for future *in vivo* reprogramming strategies. hGPCs offer an advantage to interneuron reprogramming due to their developmental origin in the ventral telencephalon^26^ and more accessible chromatin around interneuron genes^61^. Herein, hGPCs were transduced with a combination of transcription factors determining GABAergic fate, previously shown efficient in interneuron reprogramming^30,31^. Importantly, we used a 3D spheroid culture as a new reprogramming approach, providing a more complex environment to conventional 2D culture that facilitates cell communication and viability in extended time line as well as sequencing preparation^43^. Under these conditions, the hGPCs efficiently converted into neurons throughout the whole spheroid structure and revealed biomolecular changes, as seen with OPTIR. These results highlight the suitability of the 3D spheroid for efficient reprogramming and the viability of OPTIR for assessing cell composition changes in neuronal reprogramming.

Importantly, in this 3D setting we observed an efficient and immediate hGPC reprogramming into GABAergic interneurons, with a robust and rapid induction of PV gene and protein expression as early as 7 days post-transduction. Transcriptomic analysis further indicated the presence of chandelier subclass-specific characteristics and a pallidal regional identity. While pallial interneurons have been previously generated from stem cells *in vitro* and using direct reprogramming approaches in the mouse brain^14–20^, few studies to our knowledge have generated human interneurons of PV subtype with subclass specificity and elaborate neuronal morphology *in vitro*^30^. Chandelier PV subclass could represent a promising target for future therapeutic interventions, as it strongly influences the generation of gamma oscillations in the cortex and controls the firing rate of pyramidal cells^62^. Notably, the number of PV chandelier cells are decreased across several cortical areas in autism spectrum disorders and are morphologically and functionally altered in schizophrenia, epilepsy, and bipolar disorder.

Single nuclei RNA sequencing data herein revealed a rapid dynamic process of the glia-to-neuron conversion, with changes detectable as early as 3 days post-transduction when a small transcriptionally distinct cluster emerged compared to the starting population. Already by day 7, there were significant changes in gene expression profiles, marked by mature neuronal markers. Importantly, the glia-to-neuron conversion was dependent on ADLSF transduction and not due to spontaneous glia-to-neuron differentiation as confirmed by the GM and ND controls.

Induced PV neuronal cells were herein uniquely analyzed for their full transcriptomic profile as a first comprehensive transcriptomic data of reprogrammed human PV interneurons. When comparing this to previously published dataset of cortical fetal and adult human interneurons, we found a high similarity between the induced and authentic PV chandelier cells. Specifically, the gene expression patterns of our best-matching cells showed as much as 80% similarity to the endogenous PV chandelier cells. This level of resemblance is similar to previous comparisons of stem cell-derived SST neurons^63,64^ and showcases the ability of our reprogramming paradigm to reach a highly authentic cell product. Additionally, this study confirms the attainment of a mature state within only 2 weeks using ectopic expression of transcription factors, as opposed to months required for stem cell differentiation or *in vivo* development^9–11^.

To understand the gene dynamics during the neuronal conversion, we further assessed the transcriptomic shift from glia-to-PV interneurons as a first attempt using hGPCs. Our trajectory analysis revealed two distinct routes of hGPCs upon initiation of the reprogramming process; one route led to transcriptionally mature *bona fide* PV interneurons, while the other resulted in unsuccessful PV interneuron reprogramming, with cells remaining in an immature GABAergic state. The presence of two distinct routes might indicate a heterogeneity in the system, possibly influenced by the stochastic uptake of each transcription factors and their expression levels or the reprogramming potential of the starting cell population^65^. Nevertheless, these hGPC have previously shown similar neuronal conversion rates in 3D regardless of the starting glial types, suggesting that none of the glial clusters within hGPC population is inherently more prone to reprogramming^43^.

Importantly, we herein presented dynamic expression of novel genes in PV fate acquisition, whose relevance are completely new in the context of reprogramming. We observed transient upregulation of neuronal migration genes such as *PLEKHG1* and *WWOX,* possibly reflecting a more accessible chromatin around these genes, as hGPCs are very migratory cells.

*PLEKHG1*, a Rho guanine nucleotide exchange factor, known to interact with *Cdc42*^66^, is necessary for proper cortical migration of MGE progenitors^67^ and have previously been implicated in neurogenesis and neuronal migration^68^. Remarkably, *WWOX* emerged as an intermediate gene in this trajectory, a factor highly expressed in basket PV cells^69^ whose ablation leads to a significant hippocampal reduction of PV interneurons ^70^, or impairments in neuronal differentiation and migration^71^.

In addition, postsynaptic proteins such as *GPHN* and *NLGN1* were transiently upregulated. *GPHN* is a puncta marker for inhibitory synapses and is often regarded as a master regulator, essential for chandelier-mediated GABAergic transmission^62^. On the other hand, *NLGN1* is expressed at excitatory synapses, where it supports NMDA receptors and plays an important role in excitatory transmission^72^. This observed upregulation suggests that reprogrammed interneurons begin to express the necessary machinery for synaptic communication to be able to receive both excitatory and inhibitory signals. Also, markers of mitochondria biogenesis and metabolism such as *PPARGCA1*, *ESRRG*, and *RORA* were transiently upregulated. These genes are believed to orchestrate a transcriptional network crucial for PV fate by regulating mitochondria biogenesis and energy metabolism^55,73^ and highlights the metabolic support required for PV interneurons, given their high energy demands. Altogether, this suggests that interneuron reprogramming involves different developmental stages for the cells to obtain the necessary synaptic machinery and communication, as well as adjustments for energy demand^74^, to acquire a complete PV neuronal fate.

As one example, *RORA* has been implicated to orchestrate a transcriptional network supporting the metabolic need of PV interneurons^55^. In our protocol, this gene also seemed to play a functional role in the reprogramming process, and its addition showed a trend toward increased *PVALB* gene expression and increased neuronal complexity, suggesting that extrinsically delivered gene can potentially affect both fate specification and maturation during the reprogramming process. More studies are needed to fully evaluate the effects of this gene on reprogramming, but it has high potential to influence not only PV reprogramming but also stem-cell differentiation strategies. Our data also supports a transient expression of neural precursor genes in the reprogramming pathway to interneurons, which has been previously observed ^65^ and that should be further evaluated in the context of PV reprogramming and stem cell differentiation strategies.

The electrophysiological data herein showed that converted glia cells in 3D culture underwent functional maturation, improving both passive and active properties. This was further supported by the single nuclei sequencing data, which revealed upregulation of genes encoding membrane receptors that control synaptic function and voltage-gated sodium and potassium channels, such as *SCN1A* and *CACNA1A*. Nevertheless, sodium and potassium currents recorded in these cells at day 50 remained small compared to those recorded from human stem cell-derived GABAergic neurons, suggesting that the cells are still functionally immature and may not yet possess the full repertoire of membrane channels required for the fast-spiking behavior classically observed in PV interneurons. Indeed, a high density of sodium channels in the axon and potassium channels with a high activation threshold and fast deactivation are crucial for the temporal precision and frequency of action potential firing in PV interneurons^75^. Therefore, it is likely that such sophisticated properties can only be acquired once the cells integrate into a proper neuronal circuit.

In conclusion, we have herein established a rapid and reproducible 3D *in vitro* strategy for reprogramming hGPCs into pallial GABAergic interneurons with PV phenotype, exhibiting subclass characteristics closely resembling *bona fide* PV chandelier cells. This novel approach holds potential as a future cell replacement strategy and can be used for transplantation to the rodent brain to study further neuronal maturation. Importantly, it enables *in vivo* reprogramming of resident human glia precursor cells.

## Acknowledgement

This work was supported by several core facilities funded by Lund Stem cell center and MultiPark. We would like to thank Anna Hammarberg for her assistance with nuclei sorting, Jenny Johansson for cDNA library preparation, Emanuella Monni for her assistance with confocal imaging and Petter Storm for valuable suggestions and discussion. The research received funding from the Swedish Research Council (2021-01839 and 2021-03149), Knut and Alice Wallenberg Foundation (2021-0088), Crafoord foundation (20231012), Jeansson foundation (JS2018-0103), Brain Foundation (FO2019-0195), Åhléns foundation (139208), Royal Physiographical Society and Per, Ulla Schyberg Foundation, Sweden, Olle Engkvist Foundation (213-0229), Anna-Lisa Rosenberg Foundation and Royal Physiographic Society in Lund (43202). This work was also supported by Knut and Alice Wallenberg Foundation to SciLifeLab for research in Data-driven Life Science, DDLS (KAW 2020.0239) and partially supported by the Wallenberg AI, Autonomous Systems and Software Program (WASP) funded by the Knut and Alice Wallenberg Foundation.

## Author contribution

CAS: investigation; formal analysis; methodology; validation; visualization; conceptualization: writing – original draft. AD: data curation; formal analysis; investigation; methodology; validation; visualization; writing – review & editing. ECP: data curation; investigation; formal analysis; validation; visualization; writing – review & editing. AB: resources; investigation; formal analysis; writing – review & editing. JG: resources; investigation; formal analysis; visualization; writing – review & editing. EA: data curation; formal analysis; validation; supervision. ADV: formal analysis; investigation, visualization. OK: formal analysis; methodology; visualization. VO: data curation; formal analysis; methodology; supervision; validation; visualization; writing – review & editing. MP: supervision; writing – review & editing. DRO: Conceptualization; funding acquisition; project administration; visualization; resources; supervision; writing – original draft.

## Data and code availability

Processed RNA sequencing data and code are available upon request to

## References

1. Kubota, Y. (2014). Untangling GABAergic wiring in the cortical microcircuit. Curr Opin Neurobiol 26, 7–14. 10.1016/j.conb.2013.10.003.

2. Rupert, D.D., and Shea, S.D. (2022). Parvalbumin-Positive Interneurons Regulate Cortical Sensory Plasticity in Adulthood and Development Through Shared Mechanisms. Front Neural Circuits 16, 886629. 10.3389/fncir.2022.886629.

3. Hijazi, S., Smit, A.B., and van Kesteren, R.E. (2023). Fast-spiking parvalbumin-positive interneurons in brain physiology and Alzheimer’s disease. Mol Psychiatr. 10.1038/s41380-023-02168-y.

4. Arroyo-García, L.E., Isla, A.G., Andrade-Talavera, Y., Balleza-Tapia, H., Loera-Valencia, R., Alvarez-Jimenez, L., Pizzirusso, G., Tambaro, S., Nilsson, P., and Fisahn, A. (2021). Impaired spike-gamma coupling of area CA3 fast-spiking interneurons as the earliest functional impairment in the mouse model of Alzheimer’s disease. Mol Psychiatr 26, 5557–5567. 10.1038/s41380-021-01257-0.

5. Martinez-Losa, M., Tracy, T.E., Ma, K., Verret, L., Clemente-Perez, A., Khan, A.S., Cobos, I., Ho, K., Gan, L., Mucke, L., et al. (2018). Nav1.1-Overexpressing Interneuron Transplants Restore Brain Rhythms and Cognition in a MouseModel of Alzheimer’s Disease. Neuron 98, 75-+. 10.1016/j.neuron.2018.02.029.

6. Marín, O. (2012). Interneuron dysfunction in psychiatric disorders. Nat Rev Neurosci 13, 107–120. 10.1038/nrn3155.

7. Southwell, D.G., Nicholas, C.R., Basbaum, A.I., Stryker, M.P., Kriegstein, A.R., Rubenstein, J.L., and Alvarez-Buylla, A. (2014). Interneurons from Embryonic Development to Cell-Based Therapy. Science 344, 167-+. ARTN 1240622 10.1126/science.1240622.

8. Verret, L., Mann, E.O., Hang, G.B., Barth, A.M.I., Cobos, I., Ho, K., Devidze, N., Masliah, E., Kreitzer, A.C., Mody, I., et al. (2012). Inhibitory Interneuron Deficit Links Altered Network Activity and Cognitive Dysfunction in Alzheimer Model. Cell 149, 708–721. 10.1016/j.cell.2012.02.046.

9. Bershteyn, M., Bröer, S., Parekh, M., Maury, Y., Havlicek, S., Kriks, S., Fuentealba, L., Lee, S.N., Zhou, R.B., Subramanyam, G., et al. (2023). Human pallial MGE-type GABAergic interneuron cell therapy for chronic focal epilepsy. Cell Stem Cell 30, 1331-+. 10.1016/j.stem.2023.08.013.

10. Nicholas, C.R., Chen, J.D., Tang, Y.S., Southwell, D.G., Chalmers, N., Vogt, D., Arnold, C.M., Chen, Y.J.J., Stanley, E.G., Elefanty, A.G., et al. (2013). Functional Maturation of hPSC-Derived Forebrain Interneurons Requires an Extended Timeline and Mimics Human Neural Development. Cell Stem Cell 12, 573–586. 10.1016/j.stem.2013.04.005.

11. Maroof, A.M., Keros, S., Tyson, J.A., Ying, S.W., Ganat, Y.M., Merkle, F.T., Liu, B., Goulburn, A., Stanley, E.G., Elefanty, A.G., et al. (2013). Directed Differentiation and Functional Maturation of Cortical Interneurons from Human Embryonic Stem Cells. Cell Stem Cell 12, 559–572. 10.1016/j.stem.2013.04.008.

12. Vignoles, R., Lentini, C., d’Orange, M., and Heinrich, C. (2019). Direct Lineage Reprogramming for Brain Repair: Breakthroughs and Challenges. Trends Mol Med 25, 897–914. 10.1016/j.molmed.2019.06.006.

13. Pereira, M., Birtele, M., and Ottosson, D.R. (2019). Direct reprogramming into interneurons: potential for brain repair. Cell Mol Life Sci 76, 3953–3967. 10.1007/s00018-019-03193-3.

14. Torper, O., Ottosson, D.R., Pereira, M., Lau, S., Cardoso, T., Grealish, S., and Parmar, M. (2015). In Vivo Reprogramming of Striatal NG2 Glia into Functional Neurons that Integrate into Local Host Circuitry. Cell Rep 12, 474–481. 10.1016/j.celrep.2015.06.040.

15. Lentini, C., d’Orange, M., Marichal, N., Trottmann, M.M., Vignoles, R., Foucault, L., Verrier, C., Massera, C., Raineteau, O., Conzelmann, K.K., et al. (2021). Reprogramming reactive glia into interneurons reduces chronic seizure activity in a mouse model of mesial temporal lobe epilepsy. Cell Stem Cell 28, 2104–2121 e2110. 10.1016/j.stem.2021.09.002.

16. Guo, Z.Y., Zhang, L., Wu, Z., Chen, Y.C., Wang, F., and Chen, G. (2014). In Vivo Direct Reprogramming of Reactive Glial Cells into Functional Neurons after Brain Injury and in an Alzheimer’s Disease Model. Cell Stem Cell 14, 188–202. 10.1016/j.stem.2013.12.001.

17. Giehrl-Schwab, J., Giesert, F., Rauser, B., Lao, C.L., Hembach, S., Lefort, S., Ibarra, I.L., Koupourtidou, C., Luecken, M.D., Truong, D.J.J., et al. (2022). Parkinson’s disease motor symptoms rescue by CRISPRa-reprogramming astrocytes into GABAergic neurons. Embo Mol Med 14. ARTN e1479710.15252/emmm.202114797.

18. Mattugini, N., Bocchi, R., Scheuss, V., Russo, G.L., Torper, O., Lao, C.L., and Götz, M. (2019). Inducing Different Neuronal Subtypes from Astrocytes in the Injured Mouse Cerebral Cortex. Neuron 103, 1086-+. 10.1016/j.neuron.2019.08.009.

19. Heinrich, C., Bergami, M., Gascón, S., Lepier, A., Viganò, F., Dimou, L., Sutor, B., Berninger, B., and Götz, M. (2014). -Mediated Conversion of NG2 Glia into Induced Neurons in the Injured Adult Cerebral Cortex. Stem Cell Rep 3, 1000–1014. 10.1016/j.stemcr.2014.10.007.

20. Heinrich, C., Gascón, S., Masserdotti, G., Lepier, A., Sanchez, R., Simon-Ebert, T., Schroeder, T., Götz, M., and Berninger, B. (2011). Generation of subtype-specific neurons from postnatal astroglia of the mouse cerebral cortex. Nat Protoc 6, 214–228. 10.1038/nprot.2010.188.

21. Pereira, M., Birtele, M., Shrigley, S., Benitez, J.A., Hedlund, E., Parmar, M., and Ottosson, D.R. (2017). Direct Reprogramming of Resident NG2 Glia into Neurons with Properties of Fast-Spiking Parvalbumin-Containing Interneurons. Stem Cell Rep 9, 742–751. 10.1016/j.stemcr.2017.07.023.

22. Chouchane, M., de Farias, A.R.M., Moura, D.M.D., Hilscher, M.M., Schroeder, T., Leao, R.N., and Costa, M.R. (2017). Lineage Reprogramming of Astroglial Cells from Different Origins into Distinct Neuronal Subtypes. Stem Cell Rep 9, 162–176. 10.1016/j.stemcr.2017.05.009.

23. Nicolás Marichal, S.P., Ana Beltran Arranz, Chiara Galante, Franciele Franco Scarante, Rebecca Wiffen, Carol Schuurmans, Marisa Karow, Sergio Gascón, Benedikt Berninger (2023). Reprogramming early cortical astroglia into neurons with hallmarks of fast-spiking parvalbumin-positive interneurons by phospho-site deficient Ascl1. bioRxiv. 10.1101/2023.11.03.565289.

24. Levine, J.M., Reynolds, R., and Fawcett, J.W. (2001). The oligodendrocyte precursor cell in health and disease. Trends Neurosci 24, 39–47. Doi 10.1016/S0166-2236(00)01691-X.

25. Hughes, E.G., Kang, S.H., Fukaya, M., and Bergles, D.E. (2013). Oligodendrocyte progenitors balance growth with self-repulsion to achieve homeostasis in the adult brain. Nat Neurosci 16, 668-+. 10.1038/nn.3390.

26. Kessaris, N., Fogarty, M., Iannarelli, P., Grist, M., Wegner, M., and Richardson, W.D. (2006). Competing waves of oligodendrocytes in the forebrain and postnatal elimination of an embryonic lineage. Nat Neurosci 9, 173–179. 10.1038/nn1620.

27. van Tilborg, E., de Theije, C.G.M., van Hal, M., Wagenaar, N., de Vries, L.S., Benders, M.J., Rowitch, D.H., and Nijboer, C.H. (2018). Origin and dynamics of oligodendrocytes in the developing brain: Implications for perinatal white matter injury. Glia 66, 221–238. 10.1002/glia.23256.

28. Nolbrant, S., Giacomoni, J., Hoban, D.B., Bruzelius, A., Birtele, M., Chandler-Militello, D., Pereira, M., Ottosson, D.R., Goldman, S.A., and Parmar, M. (2020). Direct Reprogramming of Human Fetal- and Stem Cell-Derived Glial Progenitor Cells into Midbrain Dopaminergic Neurons. Stem Cell Rep 15, 869–882. 10.1016/j.stemcr.2020.08.013.

29. Wang, S., Bates, J., Li, X., Schanz, S., Chandler-Militello, D., Levine, C., Maherali, N., Studer, L., Hochedlinger, K., Windrem, M., et al. (2013). Human iPSC-derived oligodendrocyte progenitor cells can myelinate and rescue a mouse model of congenital hypomyelination. Cell Stem Cell 12, 252–264. 10.1016/j.stem.2012.12.002.

30. Giacomoni, J., Bruzelius, A., Stamouli, C.A., and Ottosson, D.R. (2020). Direct Conversion of Human Stem Cell-Derived Glial Progenitor Cells into GABAergic Interneurons. Cells-Basel 9. ARTN 245110.3390/cells9112451.

31. Colasante, G., Lignani, G., Rubio, A., Medrihan, L., Yekhlef, L., Sessa, A., Massimino, L., Giannelli, S.G., Sacchetti, S., Caiazzo, M., et al. (2015). Rapid Conversion of Fibroblasts into Functional Forebrain GABAergic Interneurons by Direct Genetic Reprogramming. Cell Stem Cell 17, 719–734. 10.1016/j.stem.2015.09.002.

32. Giacomoni, J., Bruzelius, A., Habekost, M., Kajtez, J., Ottosson, D.R., Fiorenzano, A., Storm, P., and Parmar, M. (2024). 3D model for human glia conversion into subtype-specific neurons, including dopamine neurons. Cell Reports Methods. 10.1016/j.crmeth.2024.100845.

33. Georgievska, B., Jakobsson, J., Persson, E., Ericson, C., Kirik, D., and Lundberg, C. (2004). Regulated delivery of glial cell line-derived neurotrophic factor into rat striatum, using a tetracycline-dependent lentiviral vector. Hum Gene Ther 15, 934–944. DOI 10.1089/hum.2004.15.934.

34. Zufferey, R., Nagy, D., Mandel, R.J., Naldini, L., and Trono, D. (1997). Multiply attenuated lentiviral vector achieves efficient gene delivery in vivo. Nat Biotechnol 15, 871–875. DOI 10.1038/nbt0997-871.

35. Giacomoni, J., Habekost, M., Cepeda-Prado, E., Mattsson, B., Ottosson, D.R., Parmar, M., and Kajtez, J. (2023). Protocol for optical clearing and imaging of fluorescently labeled rat brain slices. Star Protoc 4. ARTN 10204110.1016/j.xpro.2022.102041.

36. Gvazava, N., Konings, S.C., Cepeda-Prado, E., Skoryk, V., Umeano, C.H., Dong, J., Silva, I.A.N., Ottosson, D.R., Leigh, N.D., Wagner, D.E., et al. (2023). Label-Free High-Resolution Photothermal Optical Infrared Spectroscopy for Spatiotemporal Chemical Analysis in Fresh, Hydrated Living Tissues and Embryos. J Am Chem Soc 145, 24796–24808. 10.1021/jacs.3c08854.

37. Toplak, M., Read, S.T., Sandt, C., and Borondics, F. (2021). Quasar: Easy Machine Learning for Biospectroscopy. Cells-Basel 10. ARTN 230010.3390/cells10092300.

38. Sodersten, E., Toskas, K., Rraklli, V., Tiklova, K., Bjorklund, A.K., Ringner, M., Perlmann, T., and Holmberg, J. (2018). A comprehensive map coupling histone modifications with gene regulation in adult dopaminergic and serotonergic neurons. Nat Commun 9, 1226. 10.1038/s41467-018-03538-9.

39. Wolf, F.A., Angerer, P., and Theis, F.J. (2018). SCANPY: large-scale single-cell gene expression data analysis. Genome Biol 19. ARTN 1510.1186/s13059-017-1382-0.

40. Wolock, S.L., Lopez, R., and Klein, A.M. (2019). Scrublet: Computational Identification of Cell Doublets in Single-Cell Transcriptomic Data. Cell Syst 8, 281-+. 10.1016/j.cels.2018.11.005.

41. Fang, Z., Liu, X., and Peltz, G. (2023). GSEApy: a comprehensive package for performing gene set enrichment analysis in Python. Bioinformatics 39. 10.1093/bioinformatics/btac757.

42. Bergen, V., Lange, M., Peidli, S., Wolf, F.A., and Theis, F.J. (2020). Generalizing RNA velocity to transient cell states through dynamical modeling. Nat Biotechnol 38. 10.1038/s41587-020-0591-3.

43. Giacomoni, J., Bruzelius, A., Habekost, M., Kajtez, J., Rylander Ottosson, D., Fiorenzano, A., Storm, P., and Parmar, M. (In press. Online publication date 4th of September 2024). 3D Model for Human Glia Conversion into Subtype-Specific Neurons, Including Dopamine Neurons. Cell Reports Methods.

44. Liu, S.J., Nowakowski, T.J., Pollen, A.A., Lui, J.H., Horlbeck, M.A., Attenello, F.J., He, D., Weissman, J.S., Kriegstein, A.R., Diaz, A.A., et al. (2016). Single-cell analysis of long non-coding RNAs in the developing human neocortex. Genome Biol 17, 67. 10.1186/s13059-016-0932-1.

45. Cuddleston, W.H., Li, J., Fan, X., Kozenkov, A., Lalli, M., Khalique, S., Dracheva, S., Mukamel, E.A., and Breen, M.S. (2022). Cellular and genetic drivers of RNA editing variation in the human brain. Nat Commun 13, 2997. 10.1038/s41467-022-30531-0.

46. Ansell, B.R.E., Thomas, S.N., Bonelli, R., Munro, J.E., Freytag, S., and Bahlo, M. (2021). A survey of RNA editing at single-cell resolution links interneurons to schizophrenia and autism. RNA 27, 1482–1496. 10.1261/rna.078804.121.

47. Hu, H., and Jonas, P. (2014). A supercritical density of Na channels ensures fast signaling in GABAergic interneuron axons. Nat Neurosci 17, 686-+. 10.1038/nn.3678.

48. Juarez, P., and Cerdeño, V.M. (2022). Parvalbumin and parvalbumin chandelier interneurons in autism and other psychiatric disorders. Front Psychiatry 13. ARTN 91355010.3389/fpsyt.2022.913550.

49. Tasic, B., Yao, Z.Z., Graybuck, L.T., Smith, K.A., Nguyen, T.N., Bertagnolli, D., Goldy, J., Garren, E., Economo, M.N., Viswanathan, S., et al. (2018). Shared and distinct transcriptomic cell types across neocortical areas. Nature 563, 72-+. 10.1038/s41586-018-0654-5.

50. Hodge, R.D., Bakken, T.E., Miller, J.A., Smith, K.A., Barkan, E.R., Graybuck, L.T., Close, J.L., Long, B., Johansen, N., Penn, O., et al. (2019). Conserved cell types with divergent features in human versus mouse cortex. Nature 573, 61-+. 10.1038/s41586-019-1506-7.

51. Bakken, T.E., Jorstad, N.L., Hu, Q.W., Lake, B.B., Tian, W., Kalmbach, B.E., Crow, M., Hodge, R.D., Krienen, F.M., Sorensen, S.A., et al. (2021). Comparative cellular analysis of motor cortex in human, marmoset and mouse. Nature 598, 111-+. 10.1038/s41586-021-03465-8.

52. Favuzzi, E., Deogracias, R., Marques-Smith, A., Maeso, P., Jezequel, J., Exposito-Alonso, D., Balia, M., Kroon, T., Hinojosa, A.J., Maraver, E.F., et al. (2019). Distinct molecular programs regulate synapse specificity in cortical inhibitory circuits. Science 363, 413-+. 10.1126/science.aau8977.

53. Velmeshev, D., Perez, Y., Yan, Z.H., Valencia, J.E., Castaneda-Castellanos, D.R., Wang, L., Schirmer, L., Mayer, S., Wick, B., Wang, S.H., et al. (2023). Single-cell analysis of prenatal and postnatal human cortical development. Science 382, 173-+. ARTN eadf083410.1126/science.adf0834.

54. Zaitsev, A.V., Povysheva, N.V., Lewis, D.A., and Krimer, L.S. (2007). P/Q-type, but not N-type, calcium channels mediate GABA release from fast-spiking interneurons to pyramidal cells in rat prefrontal cortex. J Neurophysiol 97, 3567–3573. 10.1152/jn.01293.2006.

55. Paul, A., Crow, M., Raudales, R., He, M., Gillis, J., and Huang, Z.J. (2017). Transcriptional Architecture of Synaptic Communication Delineates GABAergic Neuron Identity. Cell 171, 522-+. 10.1016/j.cell.2017.08.032.

56. Wolf, F.A., Hamey, F.K., Plass, M., Solana, J., Dahlin, J.S., Göttgens, B., Rajewsky, N., Simon, L., and Theis, F.J. (2019). PAGA: graph abstraction reconciles clustering with trajectory inference through a topology preserving map of single cells. Genome Biol 20. ARTN 5910.1186/s13059-019-1663-x.

57. van Bruggen, D., Pohl, F., Langseth, C.M., Kukanja, P., Lee, H., Albiach, A.M., Kabbe, M., Meijer, M., Linnarsson, S., Hilscher, M.M., et al. (2022). Developmental landscape of human forebrain at a single-cell level identifies early waves of oligodendrogenesis. Dev Cell 57, 1421–1436 e1425. 10.1016/j.devcel.2022.04.016.

58. Elias, L.A.B., Potter, G.B., and Kriegstein, A.R. (2008). A time and a place for Nkx2-1 in interneuron specification and migration. Neuron 59, 679–682. 10.1016/j.neuron.2008.08.017.

59. Zhang, L., Qin, Z.H., Ricke, K.M., Cruz, S.A., Stewart, A.F.R., and Chen, H.H. (2020). Hyperactivated PTP1B phosphatase in parvalbumin neurons alters anterior cingulate inhibitory circuits and induces autism-like behaviors. Nature Communications 11. ARTN 101710.1038/s41467-020-14813-z.

60. Dehorter, N., Marichal, N., Marín, O., and Berninger, B. (2017). Tuning neural circuits by turning the interneuron knob. Curr Opin Neurobiol 42, 144–151. 10.1016/j.conb.2016.12.009.

61. Boshans, L.L., Factor, D.C., Singh, V., Liu, J., Zhao, C., Mandoiu, I., Lu, Q.R., Casaccia, P., Tesar, P.J., and Nishiyama, A. (2019). The Chromatin Environment Around Interneuron Genes in Oligodendrocyte Precursor Cells and Their Potential for Interneuron Reprograming. Front Neurosci 13, 829. 10.3389/fnins.2019.00829.

62. Gallo, N.B., Paul, A., and Van Aelst, L. (2020). Shedding Light on Chandelier Cell Development, Connectivity, and Contribution to Neural Disorders. Trends Neurosci 43, 565–580. 10.1016/j.tins.2020.05.003.

63. Allison, T., Langerman, J., Sabri, S., Otero-Garcia, M., Lund, A., Huang, J., Wei, X., Samarasinghe, R.A., Polioudakis, D., Mody, I., et al. (2021). Defining the nature of human pluripotent stem cell-derived interneurons via single-cell analysis. Stem Cell Rep 16, 2548–2564. 10.1016/j.stemcr.2021.08.006.

64. Close, J.L., Yao, Z.Z., Levi, B.P., Miller, J.A., Bakken, T.E., Menon, V., Ting, J.T., Wall, A., Krostag, A.R., Thomsen, E.R., et al. (2017). Single-Cell Profiling of an In Vitro Model of Human Interneuron Development Reveals Temporal Dynamics of Cell Type Production and Maturation (vol 93, pg 1035, 2017). Neuron 96, 949–949. 10.1016/j.neuron.2017.10.024.

65. Karow, M., Camp, J.G., Falk, S., Gerber, T., Pataskar, A., Gac-Santel, M., Kageyama, J., Brazovskaja, A., Garding, A., Fan, W., et al. (2018). Direct pericyte-to-neuron reprogramming via unfolding of a neural stem cell-like program. Nat Neurosci 21, 932–940. 10.1038/s41593-018-0168-3.

66. Reinhard, N.R., Van Der Niet, S., Chertkova, A., Postma, M., Hordijk, P.L., Gadella, T.W.J., Jr., and Goedhart, J. (2021). Identification of guanine nucleotide exchange factors that increase Cdc42 activity in primary human endothelial cells. Small GTPases 12, 226–240. 10.1080/21541248.2019.1658509.

67. Katayama, K., Imai, F., Campbell, K., Lang, R.A., Zheng, Y., and Yoshida, Y. (2013). RhoA and Cdc42 are required in pre-migratory progenitors of the medial ganglionic eminence ventricular zone for proper cortical interneuron migration. Development 140, 3139–3145. 10.1242/dev.092585.

68. O’Neill, A.C., Kyrousi, C., Klaus, J., Leventer, R.J., Kirk, E.P., Fry, A., Pilz, D.T., Morgan, T., Jenkins, Z.A., Drukker, M., et al. (2018). A Primate-Specific Isoform of Regulates Neurogenesis and Neuronal Migration. Cell Rep 25, 2729-+. 10.1016/j.celrep.2018.11.029.

69. Aldaz, C.M., and Hussain, T. (2020). WWOX Loss of Function in Neurodevelopmental and Neurodegenerative Disorders. Int J Mol Sci 21. ARTN 892210.3390/ijms21238922.

70. Hussain, T., Kil, H., Hattiangady, B., Lee, J., Kodali, M., Shuai, B., Attaluri, S., Takata, Y., Shen, J.J., Abba, M.C., et al. (2019). deletion leads to reduced GABA-ergic inhibitory interneuron numbers and activation of microglia and astrocytes in mouse hippocampus. Neurobiol Dis 121, 163–176. 10.1016/j.nbd.2018.09.026.

71. Kosla, K., Pluciennik, E., Orzechowska, M., Jedroszka, D., Baryla, I., Hammouz, R., Styczen-Binkowska, E., and Bednarek, A.K. (2019). The WWOX gene depletion impairs neural cell migration process and may disrupt brain development. Febs Open Bio 9, 206–206.

72. Südhof, T.C. (2017). Synaptic Neurexin Complexes: A Molecular Code for the Logic of Neural Circuits. Cell 171, 745–769. 10.1016/j.cell.2017.10.024.

73. Lucas, E.K., Dougherty, S.E., McMeekin, L.J., Reid, C.S., Dobrunz, L.E., West, A.B., Hablitz, J.J., and Cowell, R.M. (2014). PGC-1α Provides a Transcriptional Framework for Synchronous Neurotransmitter Release from Parvalbumin-Positive Interneurons. J Neurosci 34, 14375–14387. 10.1523/Jneurosci.1222-14.2014.

74. Gascón, S., Murenu, E., Masserdotti, G., Ortega, F., Russo, G.L., Petrik, D., Deshpande, A., Heinrich, C., Karow, M., Robertson, S.P., et al. (2016). Identification and Successful Negotiation of a Metabolic Checkpoint in Direct Neuronal Reprogramming. Cell Stem Cell 18, 396–409. 10.1016/j.stem.2015.12.003.

75. Hu, H., Gan, J., and Jonas, P. (2014). Interneurons. Fast-spiking, parvalbumin(+) GABAergic interneurons: from cellular design to microcircuit function. Science 345, 1255263. 10.1126/science.1255263.

