## Supplementary figures and legends for "Generating human parvalbumin interneurons through 3D glia reprogramming"

Figure S1

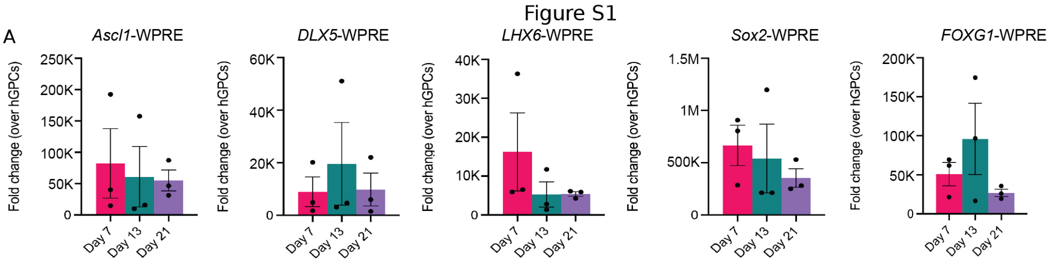
A
Ascl1-WPRE
DLX5-WPRE
LHX6-WPRE
Sox2-WPRE
FOXG1-WPRE
Fold change (over hGPCs)
250K
200K
150K
100K
50K
0
60K
40K
20K
40K
30K
20K
10K
1.5M
1M
500K
150K
50K
Day 7
Day 13
Day 21

Figure S2

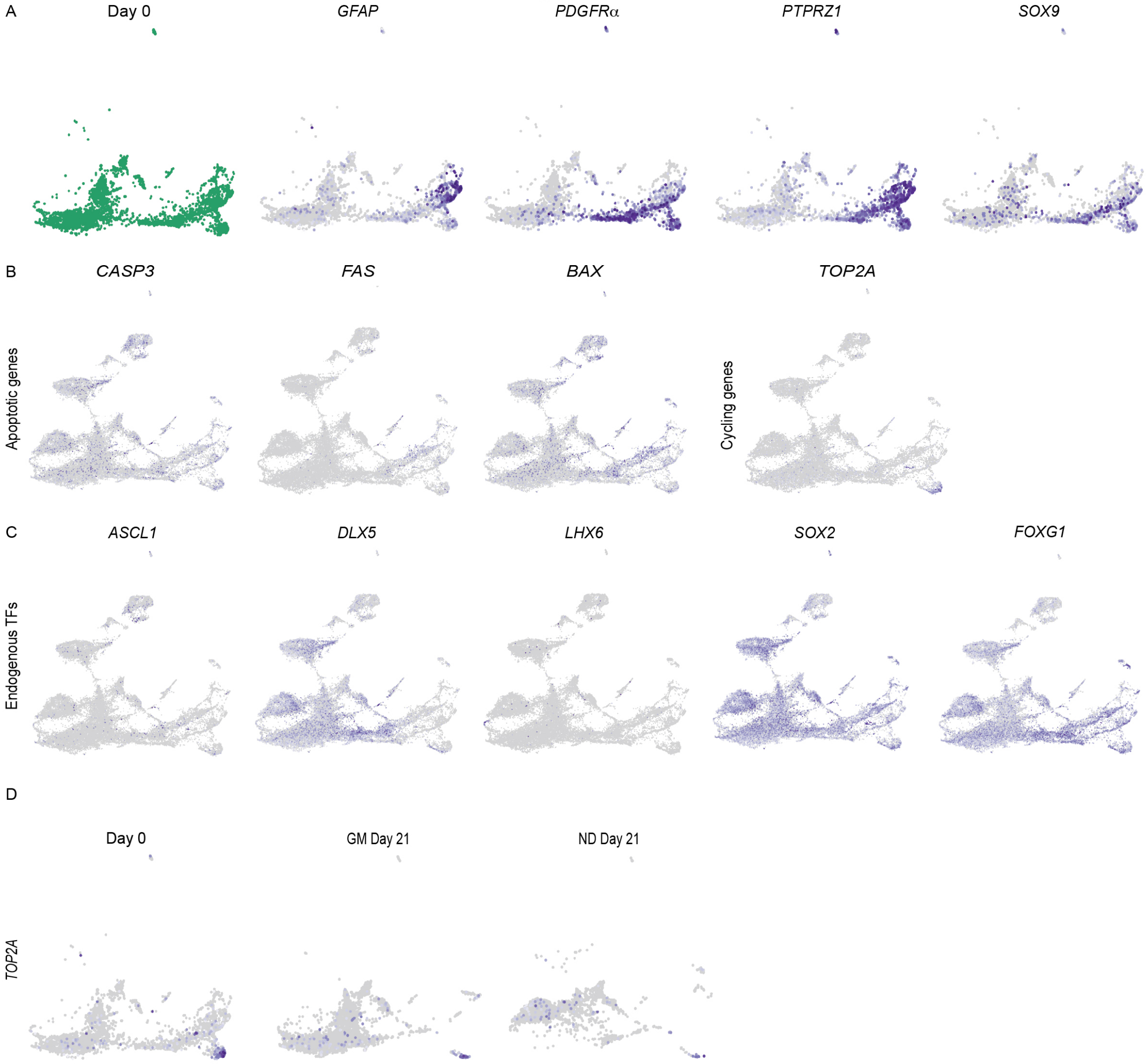
A
Day 0
GFAP
PDGFRα
PTPRZ1
SOX9
B
CASP3
FAS
BAX
TOP2A
Apoptotic genes
Cycling genes
C
ASCL1
DLX5
LHX6
SOX2
FOXG1
Endogenous TFs
D
Day 0
GM Day 21
ND Day 21
TOP2A

Figure S3

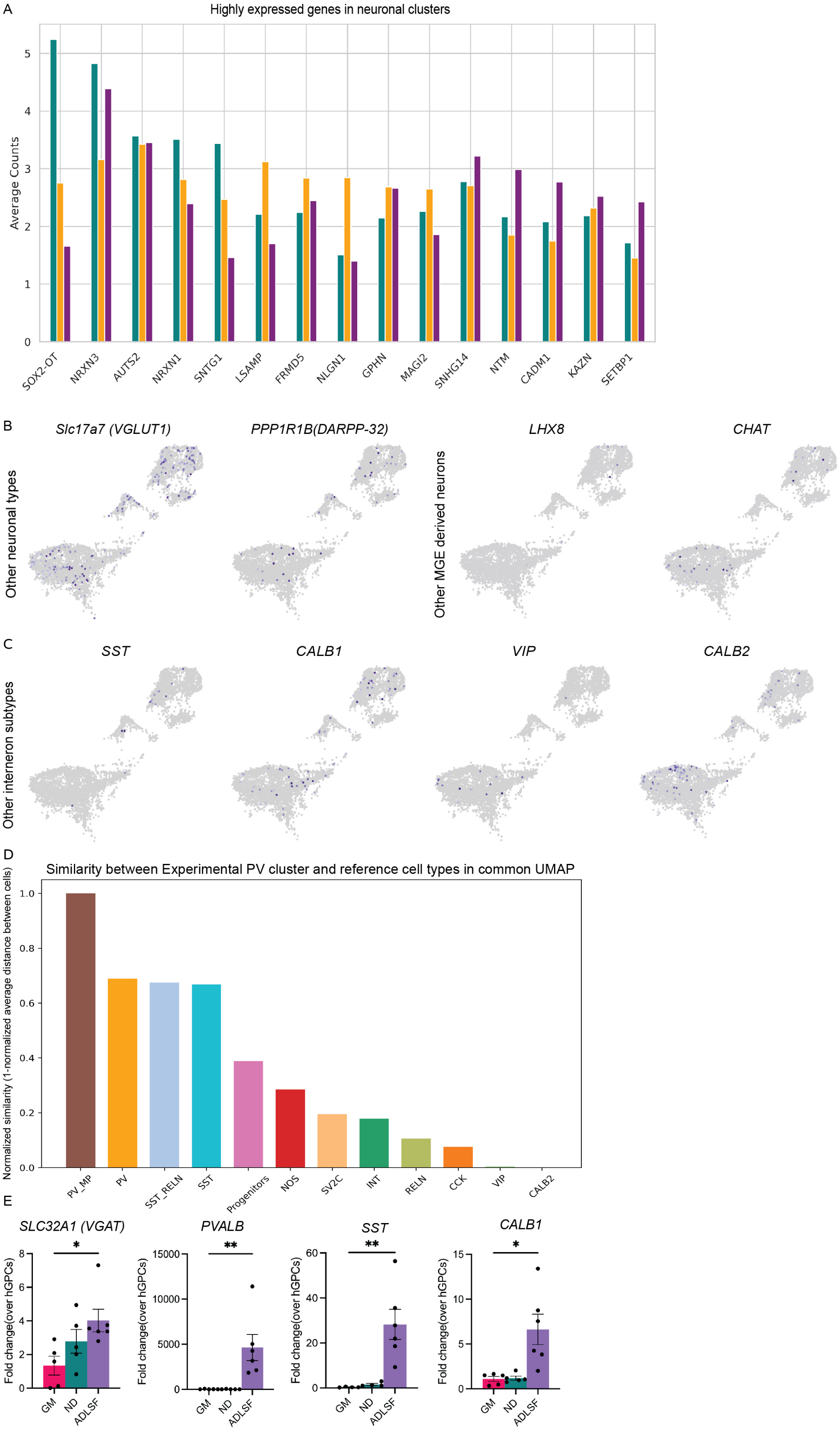

A
Highly expressed genes in neuronal clusters
Average Counts
SOX2-OT
NRXN3
AUTS2
NRXN1
SNTG1
LSAMP
FRMD5
NLGN1
GPHN
MAGI2
SNHG14
NTM
CADM1
KAZN
SETBP1
B
Slc17a7 (VGLUT1)
PPP1R1B(DARPP-32)
LHX8
CHAT
Other neuronal types
Other MGE derived neurons
C
SST
CALB1
VIP
CALB2
Other interneron subtypes
D
Similarity between Experimental PV cluster and reference cell types in common UMAP
Normalized similarity (1-normalized average distance between cells)
PV_MP
PV
SST_RELN
SST
Progenitors
NOS
SV2C
INT
RELN
CCK
VIP
CALB2
E
SLC32A1 (VGAT)
PVALB
SST
CALB1
Fold change(over hGPCs)
GM
ND
ADLSF

Figure S4

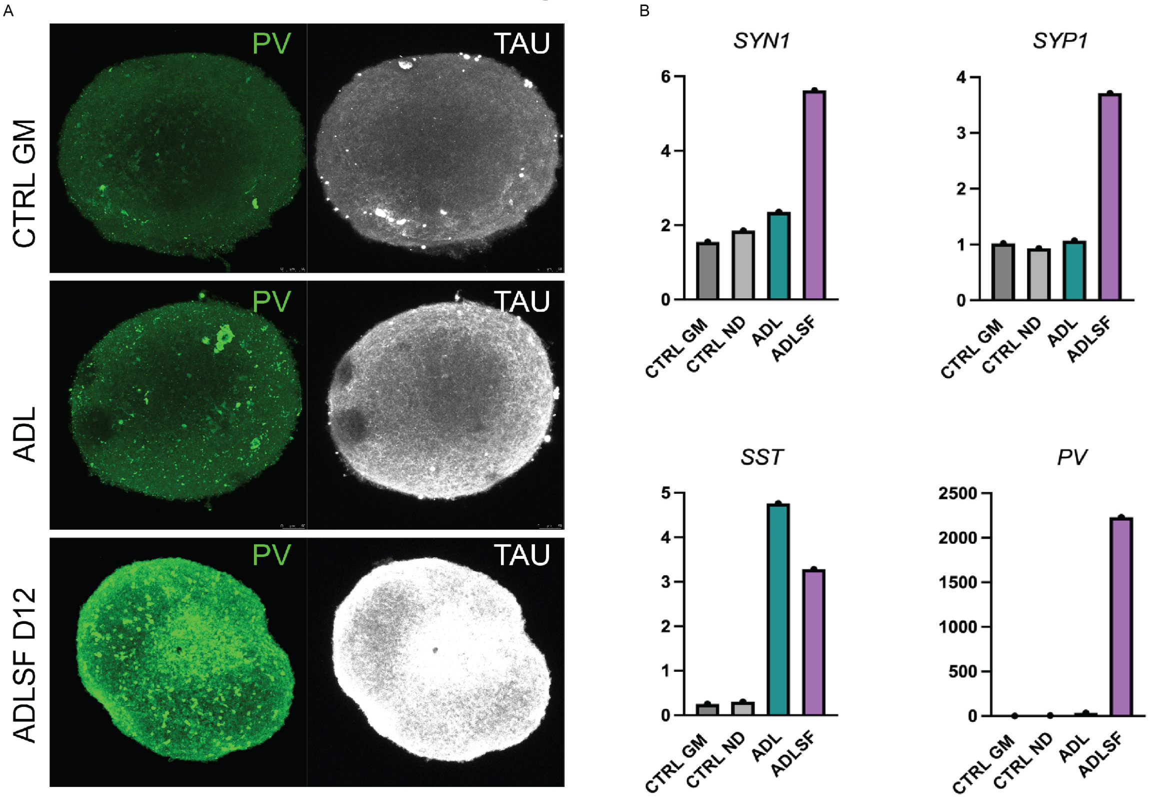
A
B
PV
TAU
CTRL GM
ADL
ADLSF D12
SYN1
SYP1
SST
PV
CTRL GM
CTRL ND
ADL
ADLSF
0
2
4
6
1
3
5
500
1000
1500
2000
2500

Figure S5

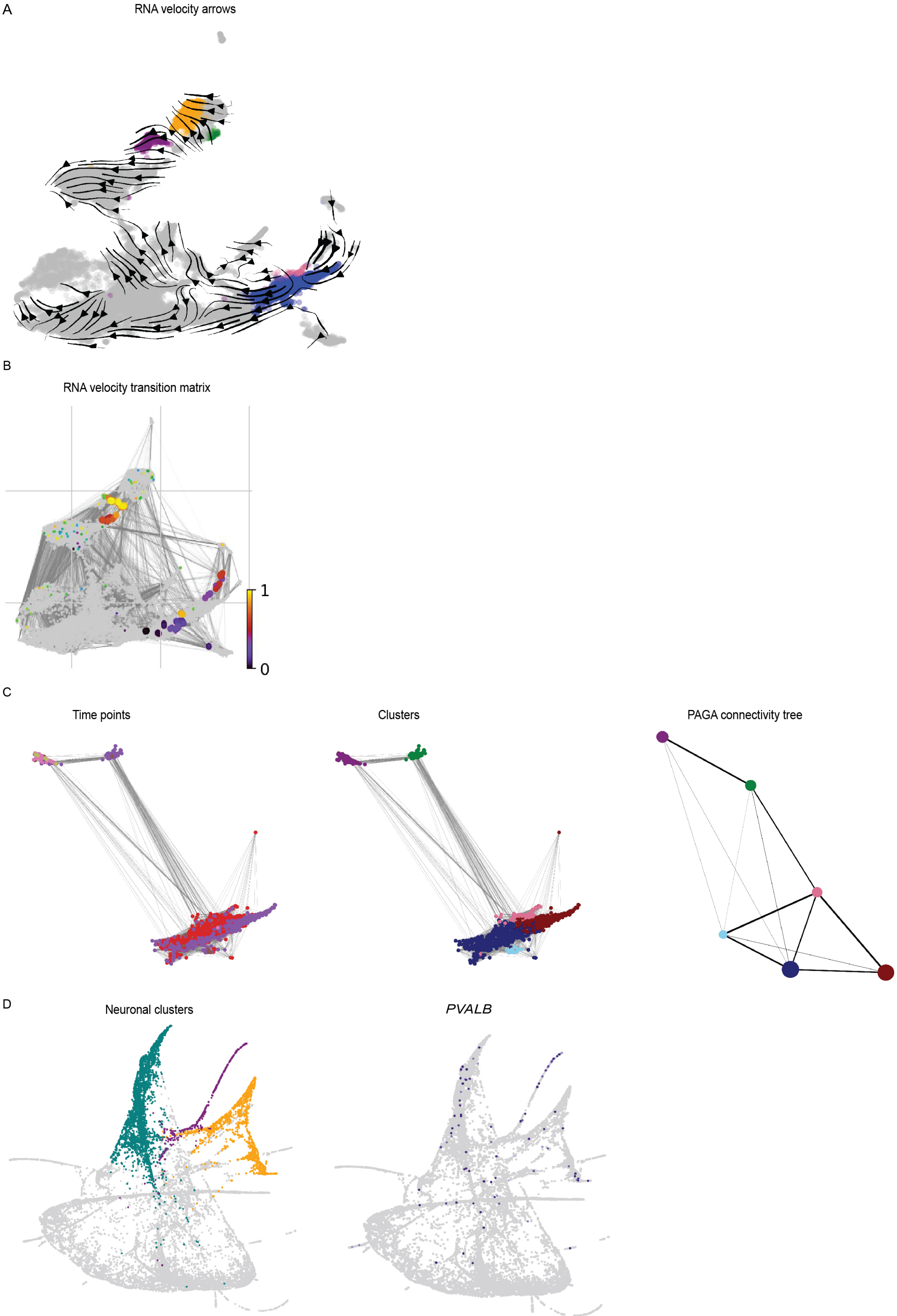
A
RNA velocity arrows
B
RNA velocity transition matrix
1
0
C
Time points
Clusters
PAGA connectivity tree
D
Neuronal clusters
PVALB

Figure S6

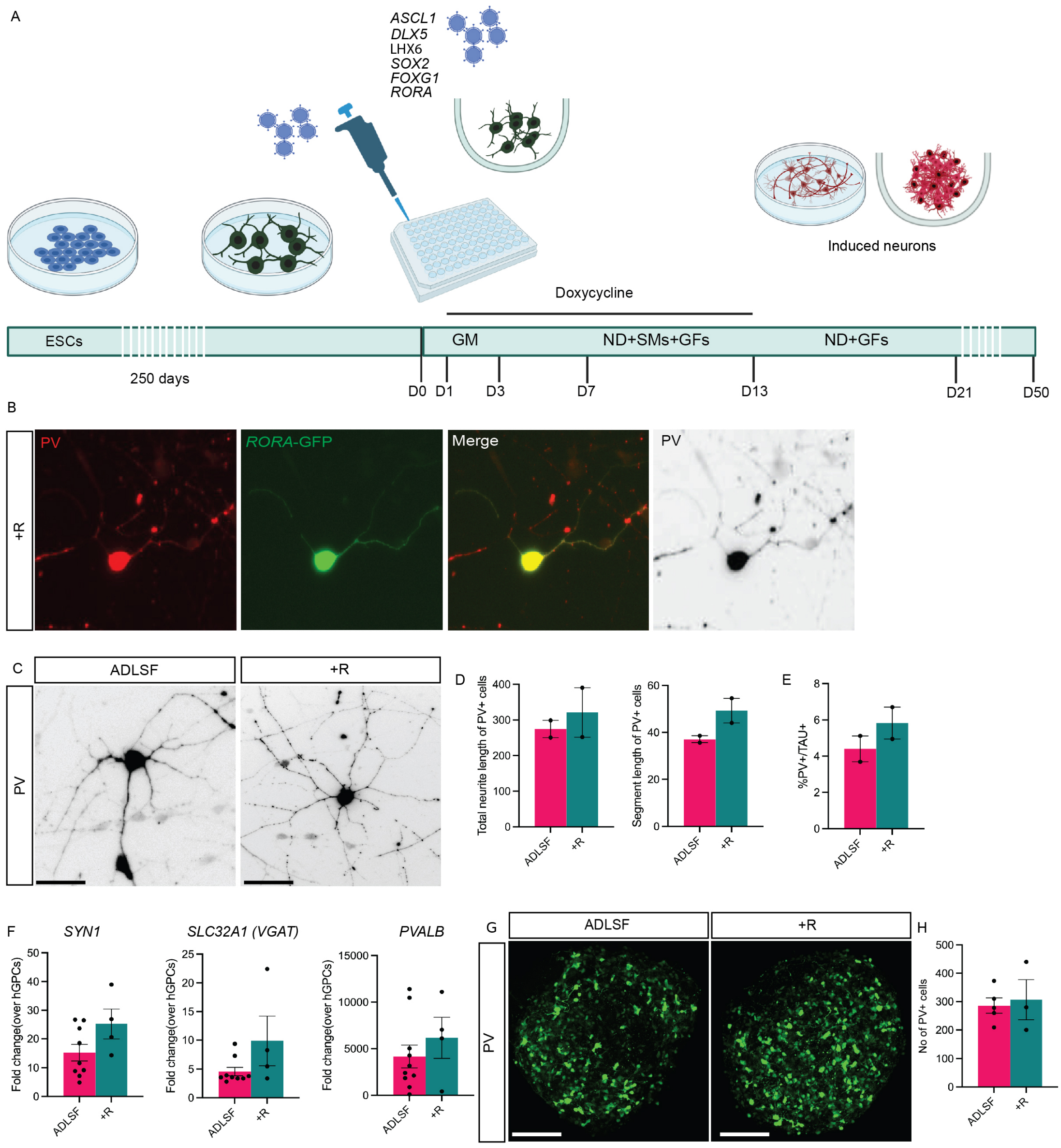
A
ASCL1
DLX5
LHX6
SOX2
FOXG1
RORA
Induced neurons
Doxycycline
ESCs
250 days
GM
ND+SMs+GFs
ND+GFs
D0
D1
D3
D7
D13
D21
D50
B
+R
PV
RORA-GFP
Merge
PV
C
ADLSF
+R
PV
D
Total neurite length of PV+ cells
Segment length of PV+ cells
ADLSF
+R
E
%PV+/TAU+
F
SYN1
SLC32A1 (VGAT)
PVALB
Fold change(over hGPCs)
G
ADLSF
+R
PV
H
No of PV+ cells

**Figure S1.** A) RT-qPCR analysis of the viral transgenes shows efficient induction of viral transcription factors (n=3 n=biological replicate). Kruskal-Wallis test was performed. Data presented as mean ± SEM.

**Figure S2.** A) UMAPs of glial markers *GFAP, PDGFRA, PTPRZ1* and *SOX9* on day 0 hGPCs (before transduction). B) UMAPs of apoptotic markers *CASP3, FAS* and *BAX* and cycling gene *TOP2A* on all the time points. C) UMAPs of all the five transcription factors, endogenous expression *ASCL1, DLX5, LHX6, SOX2, FOXG1* at all time points. D) UMAPs of cycling marker *TOP2A* on day 0 and GM and ND controls at day 21.

**Figure S3.** A) Histogram of the top highly expressed genes of the neuronal clusters. B) UMAPs of *SLC17A7* (*VGLUT1*), *PPP1R1B* (*DARPP-32*) and the cholinergic markers *LHX8* and *CHAT* on neuronal clusters. *C*) UMAPS of subtype specific markers, *SST*, *CALB1*, *VIP* and *CALB2.* D) Histogram showing the normalized similarity score between the PV-enriched cluster in this dataset and clusters in the published dataset[52], based on average distances between them in a common UMAP. E) RT-qPCR analysis of *SLC32A1(VGAT), PVALB, SST and CALB1* (n=4-6 for GM, n=4-6 for ND and n=5-6 for ADLSF, n=biological replicate) * p<0.05, **p<0.01. One way ANOVA test with post-hoc Tukey test was performed for *CALB1* and Kruskal-Wallis test with uncorrected Dunns's test for *SLC32A1*, *PVALB* and *SST.* Data presented as mean ± SEM.

**Figure S4**. A) Maximum intensity projections of immunostainings of GM control, ADL and ADLSF conditions at day 12 post conversion with PV and TAU. B) RT-qPCR analysis of *SYN1, SYP1, SST* and *PVALB* (n=1 for all conditions).

**Figure S5.** A) RNA velocity arrows on UMAP from day 1 to day 21. B) Transition matrix derived from RNA velocity showing transitions from the seed to end cells on the UMAP. C) Clusters and PAGA connectivity tree determining which sub-cluster of the starting population is most connected to the subsequent PV-trajectory clusters. D) Diffusion maps showing the neuronal clusters and expression of *PVALB*.

**Figure S6**. A) Schematic overview of the reprogramming experiments with the new factor RORA (R). B) Immunostaining of the 2D converted cells in ADLSF+R condition with PV, GFP and DAPI expression. C) Representative images PV neuron morphology for all the conditions. D) Neurite profiling of PV+/TAU+ induced neurons (n=2, n=biological replicate). E) Quantification of PV+/TAU+ induced neurons (n=2, n=biological replicate. F) RT-qPCR analysis of 3D conversions for *SYN1, VGAT* and *PVALB* (n=9-10 for ADLSF, n=4 for ADLSF+R, n=biological replicate). Unpaired t-test was performed for *SYN1* and Mann-Whitney for *VGAT* and *PVALB.* G) Maximum intensity projections of immunostainings for PV for 3D spheroids in ALDSF and ADLSF+R condition. H) Quantification of PV+ cells in ALDSF and ADLSF+R, condition (n=5 for ADLSF, n=3 for ADLSF+R, n=biological replicate). Unpaired t-test was performed. Scale bars: 100 μm. Data presented as mean ± SEM.
