## Supplementary tables for "Generating human parvalbumin interneurons through 3D glia reprogramming"

Table S1. hGPCs batches used in experiments.

| Batch ID | Days in Culture | CD140+ | CD44+ | CD140+/CD44+ |
|---|---|---|---|---|
| JAB01 | 221 | 57.0% | 42.5% | 36.8% |
| JAB04 | 205 | 52.6% | 0.5% | 1.8% |
| JAB05 | 205 | 72.5% | 1.0% | 3.5% |
| JAB04 | 172 | 43.9% | 0.5% | 0.3% |
| JAB03 | 258 | 75% | 22% | 19.2% |
| JAB02 | 270 | 83.9% | 4.7% | 7.9% |
| JAB05 | 171 | 42.7% | 13.1% | 5.5% |
| JAB04 | 206 | 60,7% | 0.2% | 2.8% |
| JAB01 | 160 | 48,9% | 0.4% | 0.6% |

Table S1. FACS analysis of the hGPCs used in experiments

Table S2. Cell ranger web summary

| Sample | No. of cells | Mean reads per cell | Median genes per cell |
|---|---|---|---|
| **Day 0** | 4,629 | 31,004 | 2,436 |
| **ADLSF Day 1** | 4,293 | 39,379 | 2,591 |
| **ADLSF Day 3** | 4,794 | 38,089 | 2,385 |
| **ADLSF Day 7** | 3,115 | 49,659 | 2,813 |
| **ADLSF Day 13** | 3,226 | 55,730 | 2,600 |
| **ADLSF Day 21** | 1,219 | 138,113 | 2,840 |
| **Ctrl GM Day 21** | 4,254 | 28,479 | 2,157 |
| **Ctrl ND Day 21** | 4,171 | 39,880 | 2,227 |

Table S2. Web summary from cell ranger for all the snRNA-seq samples

Table S3. Primers for RT-qPCR experiments.

| Gene | Gene name | Primers (Forward/Reverse) |
|---|---|---|
| ***ACTB*** | Beta-actin | CCTTGCACATGCCGGAG<br>CCTTGCACATGCCGGAG |
| ***Ascl1*-WPRE** | Achaete-Scute Family BHLH Transcription Factor 1 | CTAAAGATGCAGGTTGTGCG<br>GGAGCTTCTCGACTTCACCA |
| ***CALB1*** | Calbindin | TGGCTCCATTTCGACGCTGACG<br>ATCCAGCCTTCTTTCGCGCCTG |
| ***DLX5*-WPRE** | Distal-Less Homeobox 5-viral | GGCCTCCGGGACACTCTATT<br>AGCAGCGTATCCACATAGCG |
| ***FOXG1*-WPRE** | Forkhead Box G1-viral | AGGGGTCTTCTTCCAACCCT<br>GCAGCGTATCCACATAGCGT |
| ***GAPDH*** | Glyceraldehyde-3-Phospahte Dehydrogenase | TTGAGGTCAATGAAGGGGTC<br>GAAGGTGAAGGTCGGAGTCA |
| ***LHX6*-WPRE** | LIM Homeobox 6-viral | CGTCCACCTCAAAGCCGATA<br>GCAGCGTATCCACATAGCGT |
| ***PVALB*** | Parvalbumin | TGCAGGATGTCGATGACAGA<br>TTTCTTCAGGCCGACCATTT |
| ***SLC32A1 (vGAT)*** | Solute Carrier Family 31 Member 1 | AGATGATGAGAAACAACCCCAG<br>CACGACAAGCCCAAAATCAC |
| ***Sox2*-WPRE** | SRY-Box Transcription Factor 2-viral | TCGCACATGTGAGGATCCAA<br>AGCGTAAAAGGAGCAACATAGT |
| ***SST*** | Somatostatin | CAAGCCGCTTTAGGAGCGAG |

| | | |
|---|---|---|
| | | AGGCGGCAGGACAGCATCT |
| ***SYN1*** | Synapsin 1 | CCCGTGGTTGTGAAGATGGGGC<br>TGCCACGACACTTGCGATGTCC |
| ***SYP1*** | Synaptophysin | ACCTCGGGACTCAACACCTCGG<br>GAACCACAGGTTGCCGACCCAG |

Table S3. List of primers used in RT-qPCR experiments

Table S4. Primary antibodies for whole spheroid immunostaining

| Antibodies | Concentration | Company | Ref # |
|---|---|---|---|
| **Chicken anti-GFAP** | 1:2000 | Merck Millipore | AB5541 |
| **Goat anti-PDGFRα** | 1:300 | R&D Systems | AF-307-NA |
| **Mouse anti-TAU** | 1:500 | Thermo Fisher. Scientific | MN1000 |
| **Rabbit anti-GABA** | 1:2000 | Sigma-Aldrich | A2052 |
| **Rabbit anti-PV** | 1:500 | Swant | PV-28 |
| **Rabbit anti-Ki67** | 1:500 | Abcam | ab238020 |
| **Chicken anti-GFP** | 1:1000 | Abcam | Ab13970 |

Table S4. List of primary antibodies used in this study

Table S5. Electrophysiological parameters

| | **Days post transduction** | | | |
|---|---|---|---|---|
| | **7** | **13** | **21** | **50** |
| Resting membrane potential, mV | -34.8 ± 9.8 | -33.60 ± 3.6 | -36,72 ± 2,6 | -58.97 ± 4.0 |
| Input resistance, GΩ | 0.87 ± 0.06 | 0.72 ± 0.08 | 0.52 ± 0.08 | 0.38 ±0.08 |
| Capacitance, pF | 11.78 ± 1.9 | 14.58 ± 2.5 | 22.16 ± 2.4 | 28.65 ± 2.9 |
| Action potential amplitude, mV | 30.92 ± 0.3 | 19.58 ± 2.3 | 31,23 + 4.1 | 52.30 ± 6.6 |
| Action potential threshold, mV | -37.80 ± 0.7 | -27.64 ± 3.4 | -32.25 ± 1.4 | -38.83 ± 2.0 |
| Afterhyperpolarization, mV | -5.14 ± 3.6 | -9.00 ± 1.8 | -10.00 ± 1.8 | -16.94 ± 2.6 |

Table S5. List of electrophysiological parameters from recordings of different time points post transduction. Data presented as mean ± SEM.
